# Orchestration of pluripotent stem cell genome reactivation during mitotic exit

**DOI:** 10.1101/2023.08.09.552605

**Authors:** Silja Placzek, Ludovica Vanzan, David M. Suter

**Affiliations:** Institute of Bioengineering, Ecole Polytechnique Fédérale de Lausanne (EPFL), 1025 Lausanne, Switzerland

## Abstract

The maintenance of cell identity faces many challenges during mitosis, as most DNA-binding proteins are evicted from DNA and transcription is virtually abolished. How cells maintain their identity through cell division and faithfully re-initiate gene expression during mitotic exit is unclear. Here, we developed a novel reporter system enabling cell cycle synchronization-free separation of pluripotent stem cells in temporal bins of < 30 minutes during mitotic exit. This allowed us to quantify genome-wide reactivation of transcription, sequential changes in chromatin accessibility, and re-binding of the pluripotency transcription factors OCT4, SOX2, and NANOG (OSN). We found that transcriptional activity progressively ramped up after mitosis and that OSN rapidly reoccupied the genome during the anaphase-telophase transition. We also demonstrate transcription factor-specific, dynamic relocation patterns and a hierarchical reorganization of the OSN binding landscape governed by OCT4 and SOX2. Our study sheds light on the dynamic orchestration of transcriptional reactivation after mitosis.

## Introduction

During mitosis, chromatin is reorganized into a condensed state that disrupts the three-dimensional architecture of the genome (Gibcus et al., 2018), impairs transcription factor (TF) binding (Deluz et al., 2016; Raccaud et al., 2019) and transcriptional activity (Palozola et al., 2017). When cells exit mitosis, the whole gene regulatory landscape needs to be re-established to allow for transcriptional reactivation.

A major limitation in the field is the systematic use of cell cycle-synchronizing drugs. The vast majority of studies on mammalian gene regulation during and immediately after mitosis use nocodazole to arrest cells in prometaphase (Jordan et al., 1992; Matsui et al., 2012), followed by nocodazole wash-out to allow cells to resume progression through the cell cycle (C-S Hsiung et al., 2016; Festuccia et al., 2016; Liu et al., 2017; Palozola et al., 2017). Several hours to overnight treatment with nocodazole are used to obtain enough cells in mitosis or mitotic exit. This leads to a strong enrichment of cells in prometaphase and distorts the timing of cell cycle progression, as cells typically pass through all mitotic phases within less than 1 hour (Phillips et al., 2019). Furthermore, nocodazole treatment alters chromatin compaction and transcriptional activity beyond what can be explained by the enrichment of cells in mitosis (Shlyueva et al., 2022). Upon drug washout, individual cells display substantial variability in the timing of mitotic exit (Sarnataro et al., 2021), which blurs the dissection of genome reactivation dynamics. This raises important concerns on the physiological relevance of genome reactivation upon release from drug-mediated cell cycle synchronization.

The temporal sequence in which regulatory events involved in genome reactivation occur is mostly unknown. Promoter-enhancer contact re-establishment initiates at the anaphase to telophase transition and follows locus-specific kinetics (Zhang et al., 2019), but whether distinct TF re-binding dynamics orchestrate these changes is unclear. TF activities inferred from RNA-seq data were reported to display a TF-specific temporal ordering during mitotic exit (Sarnataro et al., 2021). However, the sequence of TF rebinding to specific sites after mitosis awaits to be dissected to shed light on the mechanistic basis of this temporal organization.

In self-renewing cells such as pluripotent stem cells (PSCs), gene regulation needs to be robustly controlled throughout the cell cycle to maintain their identity over a large number of cell divisions. The core pluripotency TFs OCT4 and SOX2 play a central role in regulating PSC self-renewal during mitotic exit (Deluz et al., 2016; Elias T. Friman et al., 2019; Liu et al., 2017). OCT4 and SOX2 form heterodimers and frequently co-occupy loci genome-wide with NANOG (Chambers and Tomlinson, 2009). The pioneer properties of OCT4 and SOX2 (Soufi et al., 2015) allow them to make chromatin accessible for the binding of non-pioneer TFs. NANOG was demonstrated to have a strong physical interaction with SOX2 and a weak one with OCT4 (Gagliardi et al., 2013; Mistri et al., 2022), suggesting distinct interaction patterns between these three TFs. How OCT4, SOX2 and NANOG orchestrate their temporal pattern of rebinding during mitotic exit remains unexplored.

Here we describe a new strategy to dissect genome reactivation during mitotic exit in unsynchronized PSCs, allowing us to decipher transcriptional reactivation dynamics and acute temporal changes in the binding of pluripotency TFs and chromatin accessibility. In combination with loss-of-function studies of each individual TF, our study reveals how TF interactions regulate their binding landscape during mitotic exit.

## Results

### Generation of a reporter system to dissect mitotic exit without drug-mediated synchronization

To overcome the limitations of drug-mediated arrest of cells in mitosis, we designed a cell cycle synchronization-free strategy to study mitotic exit under physiological conditions. Our approach is based on a fluorescent reporter system that allows us to discriminate cells at different time points just before and within 2 h after cytokinesis. Briefly, we expressed two E3 ligase substrates, the mitotic destruction domain (MD) of cyclin B1 (Kadauke et al., 2012) and hCdt1(1/100)Cy(-) (Sakaue-Sawano et al., 2017) in fusion to YPet and mCherry, respectively. This allows for the degradation of YPet and mCherry at the metaphase-anaphase transition and in S phase, respectively, in a cell cycle-controlled manner. We inserted these two constructs downstream of the PGK and EF1α constitutively active promoters, respectively, into a lentiviral vector and established a clonal mouse embryonic stem cell (mESC) line stably expressing both reporters (Cdt1-MD cell line). We then performed time-lapse fluorescence microscopy (Video S1) and tracked YPet and mCherry fluorescence intensities of individual cells during mitotic exit. YPet fluorescence decreased during late mitosis and started to slowly increase 1h after cell division. In contrast, mCherry fluorescence was high at the beginning of G1 (early G1) and was lost 2 h after cell division (late G1), in line with the activity of CUL4^Ddb1^ (Nishitani et al., 2006) (Figure 1A). The distinct temporal changes of YPet and mCherry signals resulted in characteristic cellular trajectories in a two-dimensional fluorescence space (Figure 1B and S1A), allowing to assign cells to different time windows after mitotic exit by measuring their YPet and mCherry fluorescence signals. Focusing on the first 2 h after cytokinesis, we then classified cells into four equal bins of fluorescence progression (Figure 1C). We next stained this cell line with Hoechst and determined mCherry and YPet intensities of individual cells in G1 phase by flow cytometry. The fluorescence intensities acquired by microscopy could thus be correlated to those acquired by flow cytometry. This allowed us to assign cells analyzed by flow cytometry to a given temporal bin, thus enabling us to physically separate cells at different time points after cytokinesis by FACS (Figure 1D, Figure S1B and STAR Methods). We then allocated the cells from each fluorescence bin to their respective temporal position relative to cytokinesis. We found that the four fluorescence bins corresponded to cells that advanced past cytokinesis for an average of 42, 63, 87, and 110 minutes, respectively (Figure 1E). We also reasoned that this reporter system should allow sorting for cells in anaphase and telophase. We established an enrichment strategy based on their high mCherry, intermediate YPet intensities, and high DNA content (Figure 1F). Furthermore, staining against the mitotic-specific marker H3S10p allows to select for mitotic cells (Wen et al., 2005), which we could also observe in mESCs at all stages of mitosis (Figure S1C, D).

Thus, we additionally stained our reporter cell line with an antibody against H3S10p to prevent contamination with interphase cells. Upon examination by microscopy, we found that 19.9 % of the cells were in anaphase and 12.4 % in telophase (Figure S1C-E).

**Figure 1.**
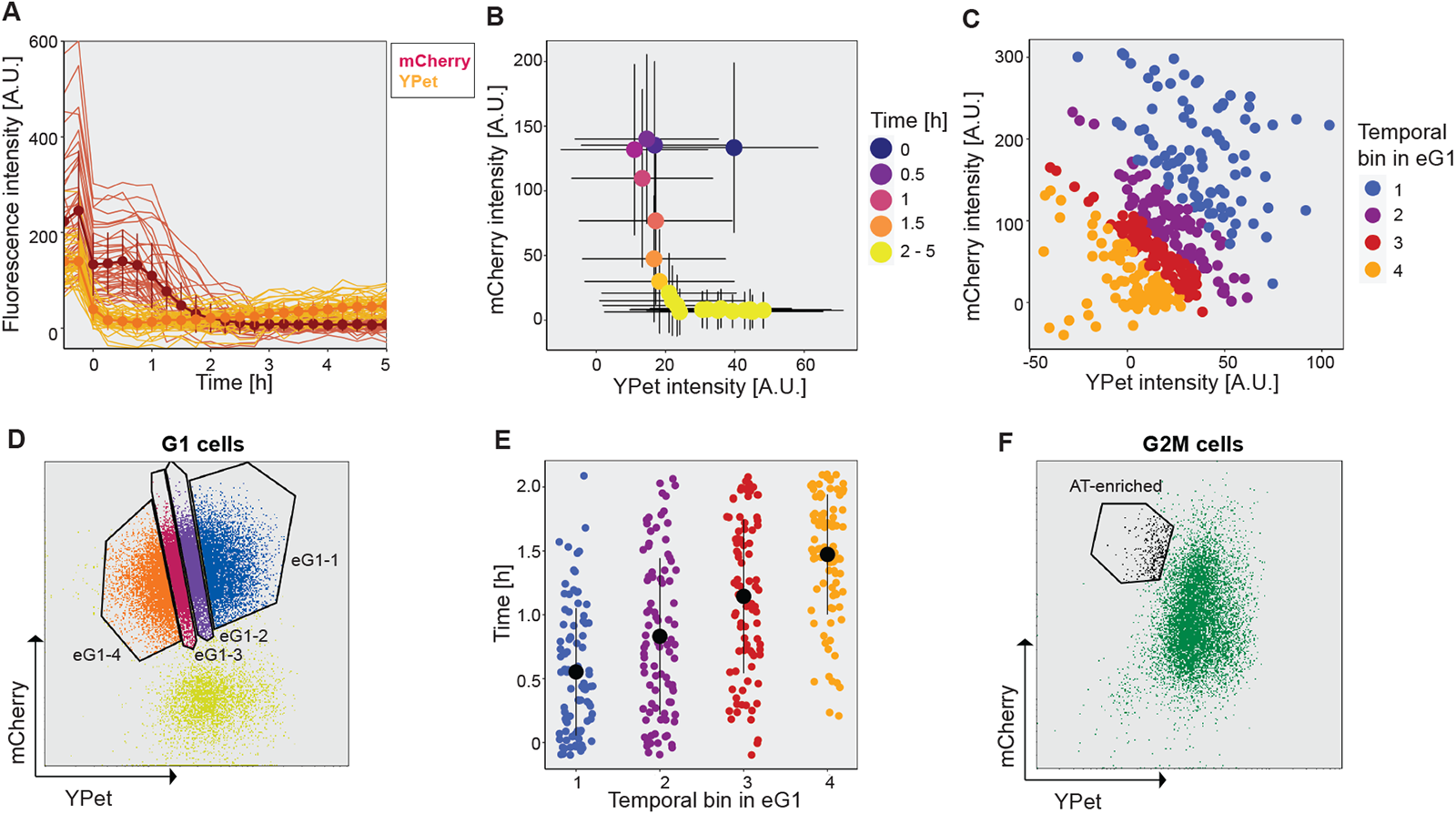
Characterization of mESCs expressing the Cdt1-MD reporter by single-cell tracking and flow cytometry analysis. (A) Temporal profile of the integrated fluorescence intensity of mCherry and YPet spanning M-G1. N=40 cells measured before cell division (time point -0.25). One sister cell was followed for 20 frames with a time interval of 15 min (5h). Dots: fluorescence mean intensity. Error bars: standard deviation (SD). Lines: single-cell traces. (B) Average integrated fluorescence intensity of mCherry and YPet for each time point of 40 cells measured over 5h after cytokinesis. Error bars: SD. (C) The first 9 time points after mitotic exit were extracted, and four temporal bins of eG1 containing an equal number of cells (as in Fig. 1D) were gated based on the reporter expression. (D) YPet and mCherry fluorescence of G1-gated cells based on Hoechst staining. Temporal bins of eG1 were gated to select an equal number of cells for each time point in eG1. Blue: eG1-1 cells; Purple: eG1-2 cells; Red: eG1-3 cells; Orange: eG1-4 cells; Yellow: late G1 cells. (E) Flow cytometry and microscopy data (Fig. 1C) were used to calculate the average time that has passed since cytokinesis in the eG1 bins. The cells left cytokinesis on average for 42 min (eG1-1, blue), 63 min (eG1-2, purple), 87 min (eG1-3, red), and 110 min (eG1-4, orange), respectively. Black dots: median; Black lines: SD. (F) YPet and mCherry fluorescence of G2M-gated cells. The gated cell population is enriched in anaphase-telophase cells, which are mCherry-high and YPet-low, and were further selected based on positive H3S10p staining.

### Transcriptional reactivation of the genome during mitotic exit

While previous studies have reported an early spike in transcriptional activity within the first hour of G1 phase, these used nocodazole-released cells blocked for several hours in prometaphase, raising the possibility that this observation may not fully reflect physiological reactivation of transcription (Chervova et al., 2023; C-S Hsiung et al., 2016; Palozola et al., 2017). We thus took advantage of the Cdt1-MD cell line to sort cells into four temporal bins of eG1 to quantify changes in transcriptional activity over time in eG1 using nuclear RNA-seq.

**Figure S1 – related to Figure 1:**
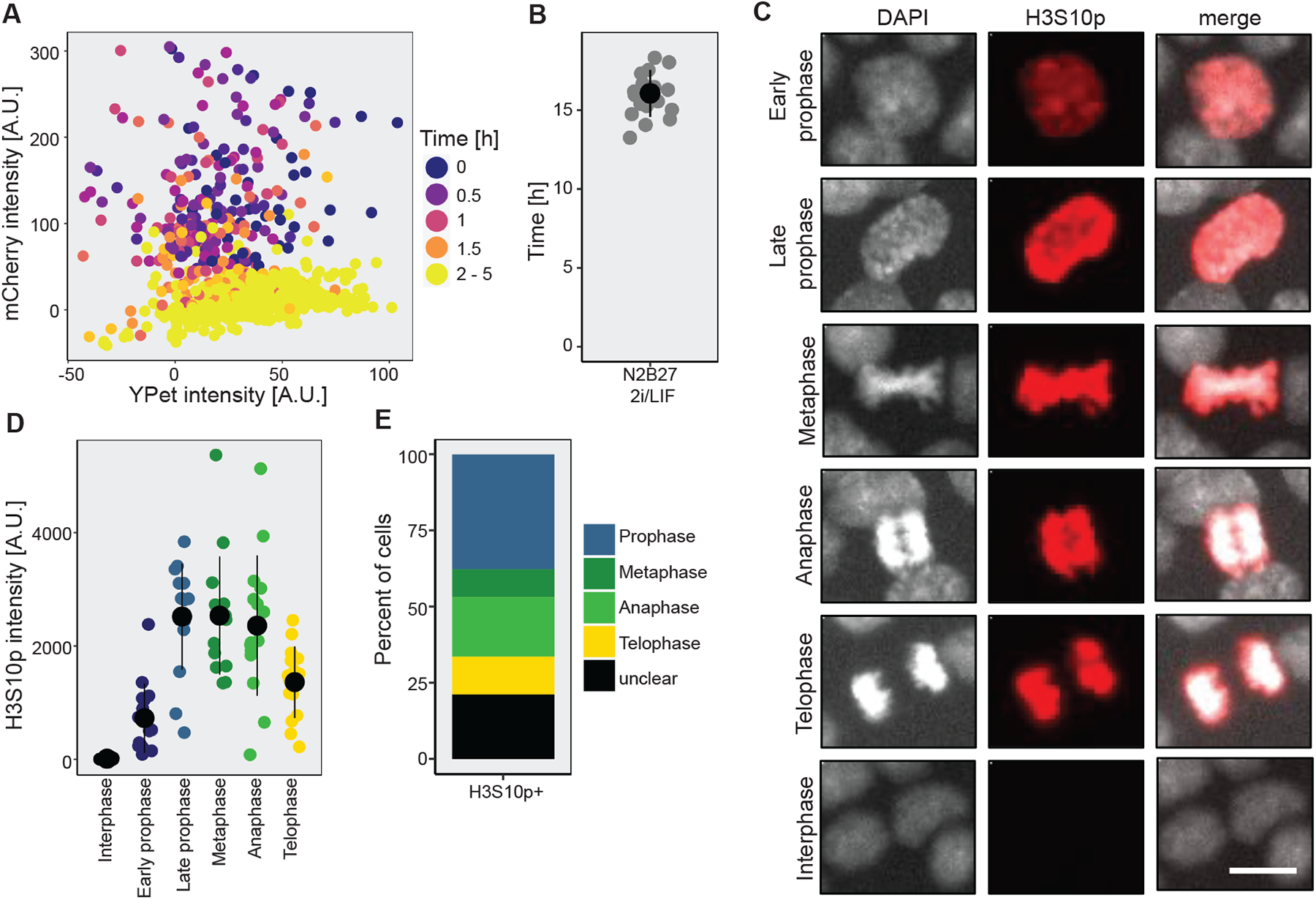
Measurements of mESCs expressing the Cdt1-MD reporter in early G1 and quantification of the H3S10p signal in mitotic cells. (A) Integrated fluorescence intensities of mCherry and YPet for each of the 20 time points of 40 cells measured over 5h after cytokinesis. (B) Duration of the cell cycle for 20 mESCs grown in N2B27 2i/LIF. Black dot: mean. Error bar: SD. (C) Representative Images and (D) immunofluorescence quantification of H3S10p signal of 15 cells per mitotic phase. Prophase was subdivided based on observed changes in chromatin condensation. Black dots: mean; Error bars: SD. Scale bar: 10 µm. (E) Cdt1-MD cells were sorted to be mCherry-high, YPet-intermediate, Hoechst-high, and H3S10p-positive, imaged, and the number of cells in each mitotic phase was counted.

In contrast to these earlier studies, we found that on average, intronic and exonic reads progressively increased and decreased, respectively, from eG1-1 to eG1-4 (Figure 2A, B). This suggests that transcriptional activity ramps up over at least 2 h after cytokinesis, while the decrease in exonic reads might be caused by a progressive increase in nuclear mRNA decay, in line with the previously reported mitosis-G1 wave of mRNA decay (Krenning et al., 2022). We then classified the dynamics of transcriptional reactivation of different genes in three clusters according to their temporal changes in unspliced (introns and intron-exon junctions) and exonic read counts (see STAR Methods). We found that most genes were already highly expressed in eG1-1 and maintained their expression at a high level throughout eG1 (Figure 2C, cluster 2). These genes are mainly involved in basic cellular processes but are also strongly enriched for the pluripotency GO term (Figure 2D). Cluster 1 genes displayed increasing unspliced and sharply decreasing exonic reads over time, suggesting transcriptional reactivation but destabilization of the mature mRNA. Cluster 3 genes displayed an opposite pattern compared to cluster 1, with a decrease in unspliced mRNAs from eG1-1 to eG1-2 but an increase in exonic reads, suggesting transcriptional shutdown and stabilization of spliced mRNAs (Figure 2C). Both clusters 1 and 3 were depleted in pluripotency terms and enriched in genes involved in cell fate commitment (Figure 2D and Table S1), suggesting that PSCs use both transcriptional and post-transcriptional mechanisms to keep the activity of genes involved in differentiation in check.

**Figure 2.**
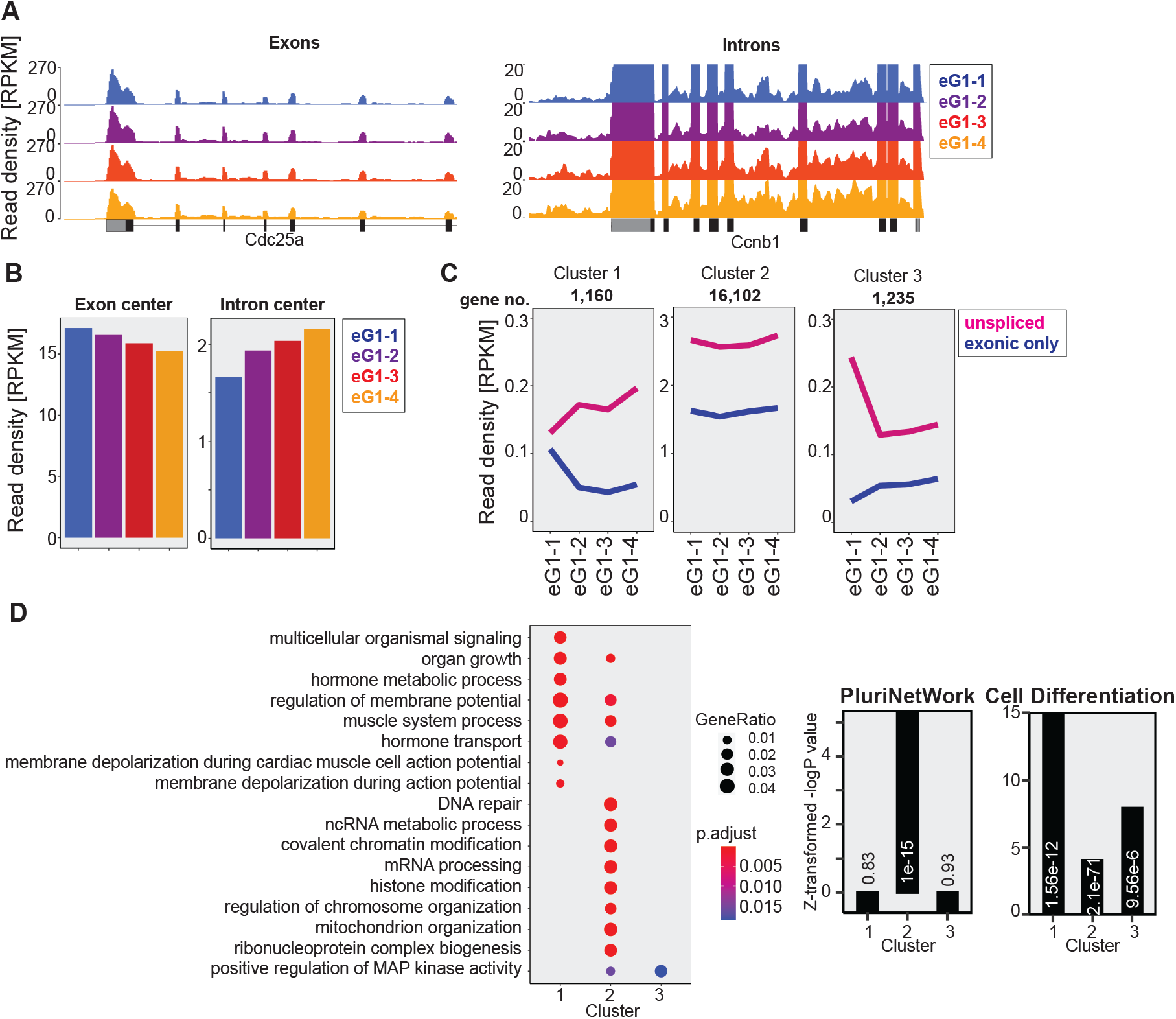
Genome reactivation of unsynchronized cells in eG1. (A) Representative examples of genome browser tracks of RNA-seq signal decreasing at exons (left) or increasing at introns (right). (B) Mean of total RNA-seq signal in different temporal bins of eG1 at the center of all exons or introns, respectively. (C) Temporal changes in read density for unspliced and exonic reads in the three clusters. The number of genes in each cluster is indicated. Lines: median. (D) GO term enrichment for the clusters shown in panel C (left) and relative enrichment values plotted as Z-transformed -logP value and p-values (digits) of GO terms PluriNetWork and Cell differentiation for the different clusters (right).

### Restoration of the chromatin landscape during mitotic exit

We then used the Cdt1-MD cell line to map genome-wide binding of the pluripotency transcription factors OCT4, SOX2, and NANOG (OSN) in unsynchronized mitotic, anaphase-telophase-enriched (AT-enriched) cells and four temporal bins of eG1 (eG1-1 to eG1-4). We sorted roughly 1 million cells for eG1-1 to eG1-4, allowing us to use 3.5 µg of chromatin for ChIP-seq. Because of the challenge to obtain large numbers of unsynchronized mitotic and AT-enriched cells, we optimized a ChIP-seq procedure using only 50,000 cells for these populations (see STAR Methods). We also acquired OSN ChIP-seq samples of eG1-1 cells with 50,000 cells to compare these three phases. This also allowed us to rescale the reads of the mitotic and AT-enriched samples to the eG1-1 to eG1-4 datasets obtained with 1 million cells, and thus to seamlessly compare all time points (see STAR Methods). All ChIP-seq data was spike-in-normalized using *Drosophila* chromatin (Bonhoure et al., 2014; Orlando et al., 2014) for quantitative comparisons between samples.

OSN bound to very few sites in mitosis (Table S2) but markedly increased their genome occupancy in AT-enriched cells to reach similar levels over the whole genome in eG1-1 (Figure 3A and Figure S2B). Importantly, we also generated an OCT4 ChIP-seq dataset from 1 million unsynchronized mitotic cells, which confirmed the very low mitotic occupancy of OCT4. This suggests that the low number of peaks found in the 50,000-cells mitotic samples cannot be simply explained by the lower chromatin input (Figure S2A). To dissect locus-specific changes in TF binding during mitosis, we next performed a Trimmed Mean of M-values (TMM)-normalization (Robinson and Oshlack, 2010) on the ChIP-seq data, which allows for normalization of technical variation during sample preparation (Figure S2C-D). We then quantified temporal changes in OSN binding to regulatory elements. We found that all three TFs were mainly occupying promoters and active enhancers before progressively spreading to other genomic regions during AT and eG1 (Figure 3B). To quantify locus-specific temporal changes in TF binding during mitotic exit, we next performed a track subtraction (see STAR Methods) between each two consecutive timepoints for each TF, and quantified changes in peak intensities at all bound loci. Genomic loci were ordered according to quantitative changes between mitosis and AT-enriched cells (Figure 3C). TF occupancy increased from mitosis to AT-enriched cells at all loci, while changes in TF occupancy were more bidirectional between later time points. Interestingly, loci that most strongly increased their binding occupancy from mitosis to AT-enriched cells tended to further increase their binding in eG1-1, and conversely for those that lost binding (Figure 3C), suggesting a progressive and consistent TF relocation from mitosis to eG1. For better visualization, we also quantified the magnitude of these changes from eG1-1 to eG1-4 in violin plots and their standard deviation, which show that OCT4 and NANOG relocalization occurred throughout eG1-1 while SOX2 was mainly redistributed between eG1-1 and eG1-2 (Figure 3C).

**Figure 3.**
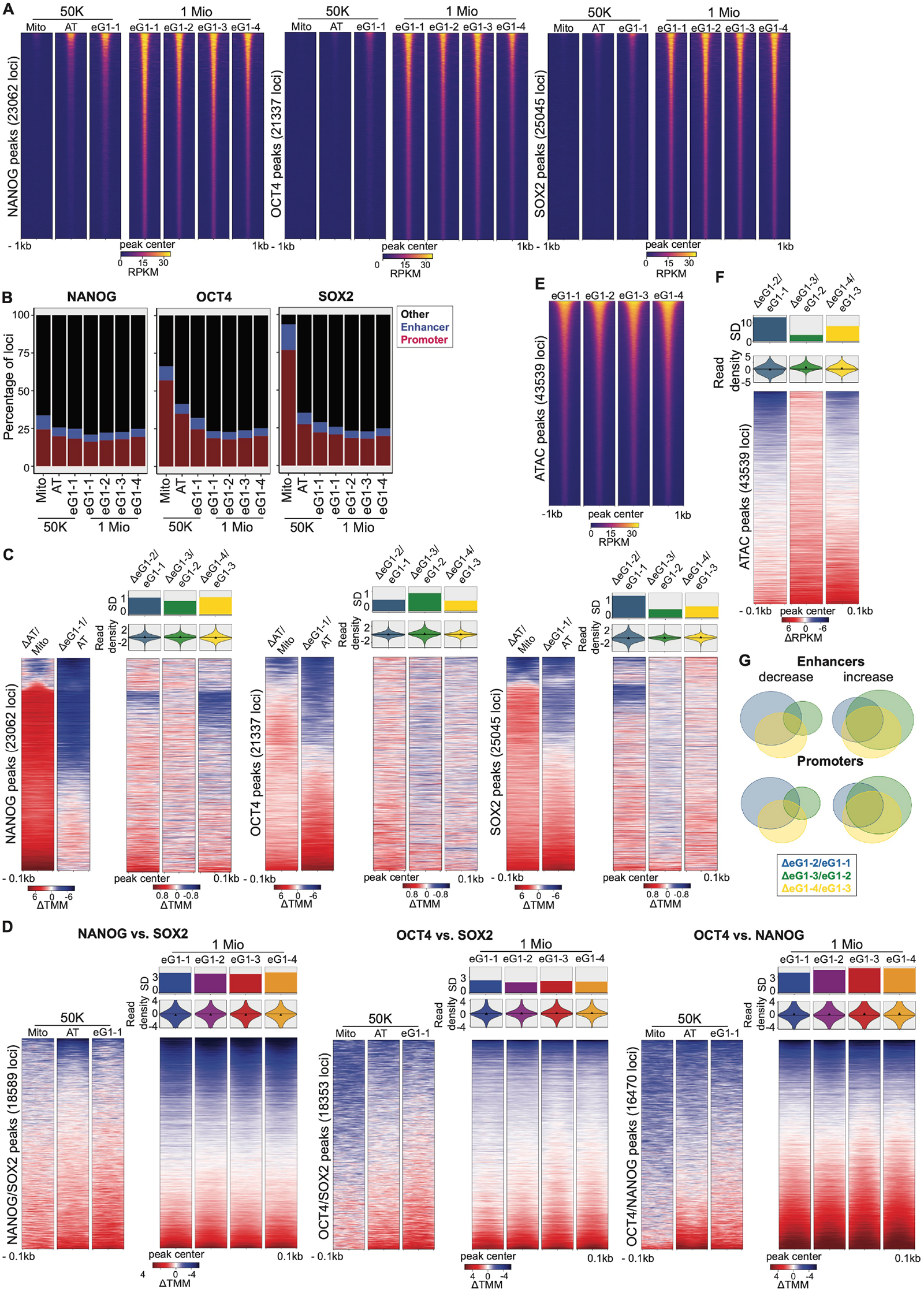
TF binding and chromatin accessibility changes throughout mitotic exit. (A) Heatmaps of spike-in and RPKM-normalized ChIP-seq binding of OSN in mitosis (Mito), AT-enriched cells (AT), and different temporal bins of eG1. Signals are ranked by intensity, and each row represents one locus. (B) Overlap of OSN binding sites with active enhancer and promoter peaks at different time points. (C) Heatmaps of ΔTMM track-subtracted reads between different timepoints for OSN on all peaks. Positive (red) and negative (blue) values: increase and decrease in signal, respectively. SD of the values plotted in the heatmaps and violin plots. Triangles: median. (D) Heatmaps of ΔTMM track-subtracted read density between the indicated TFs 100bp around their common peaks. Positive (red) and negative (blue) values: increase and decrease in overlap, respectively. SD of the values plotted in the heatmaps and violin plots. Triangles: median. (E) Heatmaps of RPKM-normalized ATAC-seq data on different temporal bins of eG1. (F) Heatmaps of the ΔRPKM track subtracted ATAC-seq reads between each subsequent timepoint of the ATAC-seq data. Positive (red) and negative (blue) values: increase and decrease in signal, respectively. SD of the values plotted in the heatmaps and violin plots. Triangles: median. (G) Overlap between active enhancers and promoters that decrease or increase in accessibility between different time points in eG1.

As OSN colocalize at many loci in the genome (Chen et al., 2008), we next asked how the co-binding behavior between OSN changes over time by performing a track subtraction between each consecutive time point for each TF pair. Using this metric on the TMM-normalized data, a perfect co-localization at a given locus will result in a value of 0, while any deviation from this value means that one TF binds more than the other relative to its genome-wide binding. As expected, we found that OCT4 and SOX2 were more co-localized together than with NANOG, in line with their binding as a heterodimer at most genomic loci (Chen et al., 2008; Rodda et al., 2005). While the co-binding of SOX2 with NANOG and OCT4 stayed constant, the co-binding of OCT4 with NANOG slightly decreased over time (Figure 3D).

Next, we quantified chromatin accessibility changes throughout eG1 using ATAC-seq. We found that global chromatin accessibility was already mostly restored in eG1-1 (Figure 3E, Figure S2E-F). We then quantified changes in accessibility of promoters and active enhancers throughout eG1. To do so, we performed a track subtraction between consecutive time points (Figure 3F) and extracted loci that increased or decreased in accessibility over time. We found that the largest changes in accessibility occurred between eG1-1 and eG1-2, and that a majority of active promoters and enhancers increased in accessibility over time (Figure S2G). The loci increasing accessibility between two consecutive time points were mostly the same throughout eG1, in contrast to loci losing accessibility (Figure 3G, Table S4). Therefore, a large fraction of regulatory regions display a consistent, gradual re-establishment of chromatin accessibility throughout eG1, while the closing of regulatory regions during eG1 is less coordinated in time.

### Post-mitotic binding dynamics of OSN display distinct relationships to local chromatin accessibility

We next classified OSN binding sites into clusters characterized by distinct dynamic changes in occupancy (see STAR Methods) and related these to local chromatin accessibility. We identified three clusters that displayed increased, stable, or decreased TF enrichment throughout eG1. For NANOG, clusters 1 and 2 displayed high chromatin accessibility, in contrast to cluster 3 that was less accessible (Figure 4A). We then quantified the overlap of these clusters with loci that depend on the pioneer activity of OCT4 and SOX2. These regions were defined by measuring changes in chromatin accessibility after acute depletion of either OCT4 or SOX2, allowing us to determine which loci depend only on the pioneering activity of OCT4 (OCT4-dependent), only on the pioneering activity of SOX2 (SOX2-dependent) or required the pioneering activity of both OCT4 and SOX2 (Co-dependent) to maintain their accessibility (Friman et al., 2019). We found that clusters 1 and 2 were more enriched in OCT4-dependent and Co-dependent regions than cluster 3 (Figure S3A). In contrast, cluster 3 was more enriched in SOX2-dependent regions (Figure S3A), less enriched in pluripotency terms, and more in differentiation terms (Figure 4A). Therefore, NANOG initially scans regions that are both less accessible and less related to pluripotency maintenance, to later relocate to pluripotency regulatory regions made accessible by the activity of OCT4 alone or OCT4 and SOX2 together. In the case of OCT4, regions increasing its enrichment from eG1-1 to eG1-4 were the most accessible, while those decreasing OCT4 binding were the least accessible. This suggests that pre-existing chromatin accessibility fosters and sustains robust OCT4 binding during eG1. Interestingly, pluripotency and differentiation terms were both most enriched in regions increasing or maintaining high OCT4 binding (Figure 4B). Finally, SOX2 binding displayed a strong preference for accessible regions during eG1-1 before relocating to less accessible regions in eG1-2 (Figure 4C). Overall, these results indicate that different pluripotency TFs exhibit distinct relocalization dynamics upon mitotic exit, associated with TF-specific chromatin accessibility profiles and functional gene categories.

**Figure S2 – related to Figure 3:**
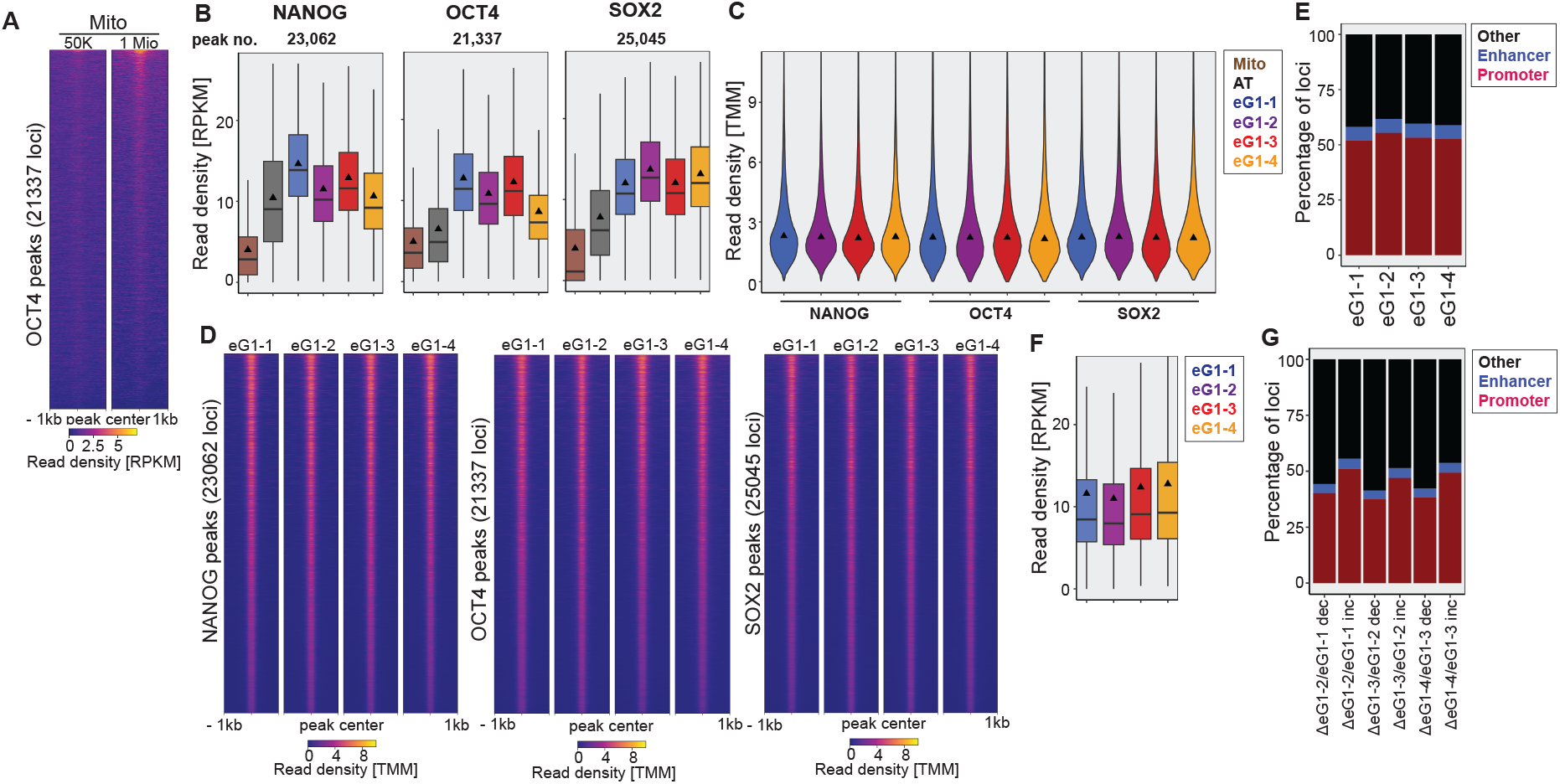
TMM-normalization of ChIP-seq data and signal quantification of track-subtracted loci. (A) Heatmap of spike-in and RPKM-normalized ChIP-seq data on OCT4 in mitosis on 50,000 or 1 Mio unsynchronized cells on all consensus peaks of OCT4 from mitosis to eG1-4. Signals are ranked by intensity, and each row represents one locus. (B) Boxplots of spike-in and RPKM-normalized ChIP-seq data on OSN in mitosis (Mito), AT-enriched cells (AT), and different temporal bins of eG1. Mitotic and AT reads were scaled based on the difference between the number of reads in eG1-1 on 1 Mio and 50,000 cells (see STAR Methods). Triangles: mean. (C) Violin plots of the TMM-normalized reads of OSN in different temporal bins of eG1 after TMM-normalization on the reference peaks. Triangles: median. (D) Heatmaps of TMM-normalized ChIP-seq data on OSN in different temporal bins of eG1 1kb around all consensus peaks of each respective TF from mitosis to eG1-4. Signals are ranked by intensity, and each row represents one locus. (E) Overlap of ATAC-seq peaks with active enhancers and promoters. (F) Boxplots of RPKM-normalized ATAC-seq data in different temporal bins of eG1. (G) Overlap of ΔRPKM track subtracted-ATAC-seq peaks with active enhancers and promoters at different time points of eG1. For boxplots: Boxes: intervals between the 25th and 75th percentile. Horizontal lines: median; Triangles: Mean; Error bars: 1.5-fold the interquartile range or the closest data point when the data point is outside this range.

**Figure 4.**
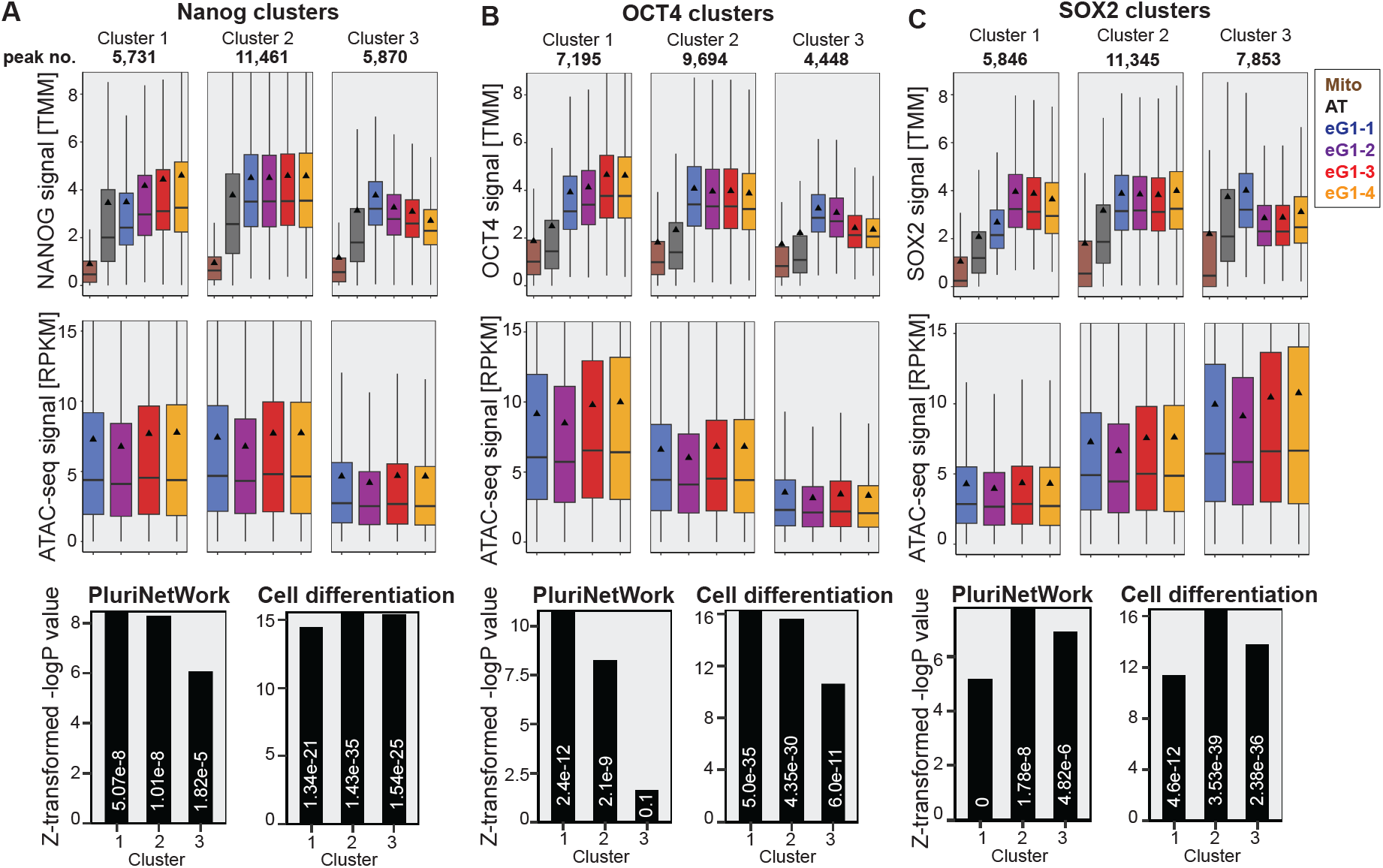
OSN are redistributed through eG1. (A-C) Boxplots of the TMM-signal after clustering of NANOG (A), OCT4 (B), or SOX2 (C) binding. Boxes: intervals between the 25th and 75th percentile. Horizontal lines: median; Triangles: mean; Error bars: 1.5-fold the interquartile range or the closest data point when the data point is outside this range. RPKM-signal of ATAC-seq data underneath. Last row: relative enrichment values of the Z-transformed -logP value and p-value (digits) of GO term enrichment of PluriNetWork and Cell differentiation.

### OCT4 and SOX2 orchestrate genome-wide NANOG occupancy during mitotic exit

To further dissect the orchestration of TF re-binding during mitotic exit, we probed how OSN binding depend on the presence of each other. To do so, we used the ZHBTc4 and 2TS22C cell lines, in which OCT4 (Niwa et al., 2000) or SOX2 (Masui et al., 2007), respectively, can be depleted using a doxycycline-inducible Tet-off system. In addition, we used the 44iN (Festuccia et al., 2012) cell line, in which NANOG is controlled by a Tet-on system and can be depleted by doxycycline removal. While OCT4 and SOX2 loss had a strong impact on the binding of the two other respective TFs and on chromatin accessibility at their binding sites, NANOG loss had only a mild impact on these (Figure 5A). We thus focused on how NANOG binding dynamics correlated with OCT4 and SOX2 binding dynamics during mitotic exit.

**Figure S3 – related to Figure 4 and 5:**
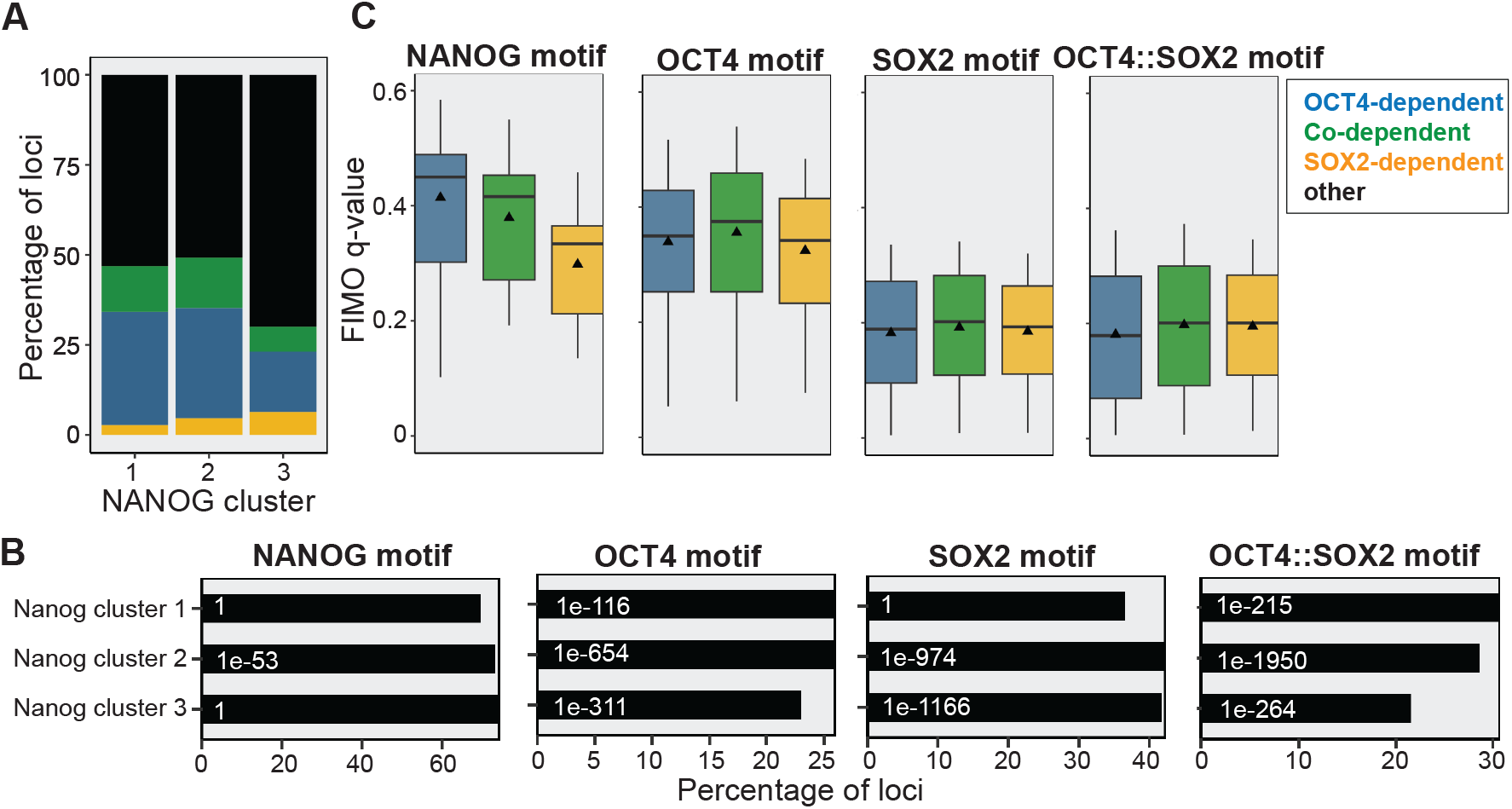
Additional characterization of OSN binding upon mitotic exit and Nanog cluster 1-3. (A) Overlap between the binding sites of the different Nanog clusters with OCT4-, SOX2-, and Co-dependent loci as defined in (Friman et al., 2019). The remaining sites were classified as other. Percentages were calculated based on the total number of peaks in each cluster. (B) NANOG, OCT4, SOX2, and OCT4::SOX2 motif enrichment and p-value (white digits) in Nanog cluster 1-3. (C) NANOG, OCT4, SOX2, and OCT4::SOX2 motif quality based on FIMO q-value in Nanog cluster 1-3. Boxes: intervals between the 25th and 75th percentile. Horizontal lines: median; Triangles: mean; Error bars: 1.5-fold the interquartile range or the closest data point when the data point is outside this range.

OCT4 and SOX2 were almost maximally enriched in eG1-1 in NANOG cluster 1, where NANOG binding increases over time (Figure 5B). Cluster 1 was also most highly enriched in the OCT4::SOX2 motif (Figure S3B-C). This is compatible with the progressive relocalization of NANOG to cluster 1 being dependent on the strong and early binding of OCT4 and SOX2 to these loci. We next used the datasets in which OCT4 (this study) or SOX2 (this study and Friman et al., 2019) were depleted to interrogate how NANOG binding changes at the three NANOG clusters. While the loss of OCT4 and SOX2 led to a decrease in NANOG enrichment at all clusters (Figure 5C-D), The impact of SOX2 loss and OCT4 loss were strongest and weakest, respectively, on both NANOG binding and chromatin accessibility in cluster 3 as compared to cluster 1 and 2. This suggests that SOX2 plays a stronger role than OCT4 for NANOG to bind in cluster 3 loci.

We then further dissected how OCT4 and SOX2 impact NANOG binding using a differential binding analysis on NANOG ChIP-seq in the presence or absence of OCT4 or SOX2. We found that 17 % and 13 % of loci decreased NANOG binding in the absence of OCT4 and SOX2, respectively, and 1.4 % of loci increased NANOG binding in the absence of OCT4 (Figure 5E-F, Table S3). We next performed ChIP-seq on SOX2 and OCT4 in the presence or absence of OCT4 or SOX2, respectively, and using a published ATAC-seq dataset acquired in the same conditions (Friman et al., 2019). Upon loss of OCT4 or SOX2, the loci that significantly decreased NANOG binding also lost the binding of SOX2 or OCT4, respectively, and became inaccessible (Figure 5E-F). Loci that significantly increased NANOG binding in the absence of OCT4 were accompanied by an increase in SOX2 binding and chromatin accessibility (Figure 5E). This indicates that in the absence of OCT4, SOX2 and NANOG are redirected to new sites that become accessible.

We then quantified how OCT4 or SOX2 depletion impact NANOG binding at loci that change NANOG binding in the absence of SOX2 or OCT4, respectively. Upon loss of either TF, sites that decreased NANOG binding in the absence of one TF also decreased NANOG binding in the absence of the other TF and became less accessible (Figure 5G-H). This indicates that both OCT4 and SOX2 are required to maintain chromatin accessibility and allow NANOG binding at these sites. We next identified the loci that increased, stayed stable, or decreased SOX2 or OCT4 binding in the absence of OCT4 or SOX2, respectively, and quantified TF binding changes and chromatin accessibility in these regions. Interestingly, loss of OCT4 resulted in an increased SOX2 binding at 1.8% of loci (Figure S4A), while very few loci increased OCT4 binding upon loss of SOX2 (Figure S4B). Changes in OCT4 and SOX2 binding were always accompanied by corresponding changes in NANOG enrichment and chromatin accessibility (Figure S4A and Figure S4B), and loci that decreased NANOG binding broadly overlapped with the ones that decreased SOX2 or OCT4 binding (Figure S4C, Table S4). Moreover, regions losing NANOG binding highly overlapped with regions depending on OCT4, SOX2 or both for their accessibility (Figure S4D) and were highly enriched in OCT4, SOX2, and OCT4::SOX2 motifs (Figure S4E-F). Regions where NANOG binding increased upon OCT4 loss had strong NANOG and SOX2 motifs but weak OCT4 and OCT4::SOX2 motifs (Figure S4G-H) and were depleted in GO terms involved in pluripotency (Figure S4I-J). Furthermore, these regions lost NANOG binding upon SOX2 loss (Figure 5G), indicating that SOX2 is required for NANOG binding to those loci.

Taken together, these results indicate that OCT4 and SOX2 broadly cooperate to facilitate NANOG binding at pluripotency regulatory elements and that in the absence of OCT4, SOX2 redirects NANOG to loci that are not involved in pluripotency maintenance. These results echo the lower and higher dependency of NANOG on the pioneering activity of OCT4 and SOX2, respectively, during eG1-1 (cluster 3 in Figure S3A), leading to its binding to loci less enriched for pluripotency terms (cluster 3 in Figure 4A), and subsequent NANOG relocation to loci that are OCT4-dependent or Co-dependent and more enriched for pluripotency terms (clusters 1 and 2 in Figure S3A and Figure 4A, respectively).

**Figure 5.**
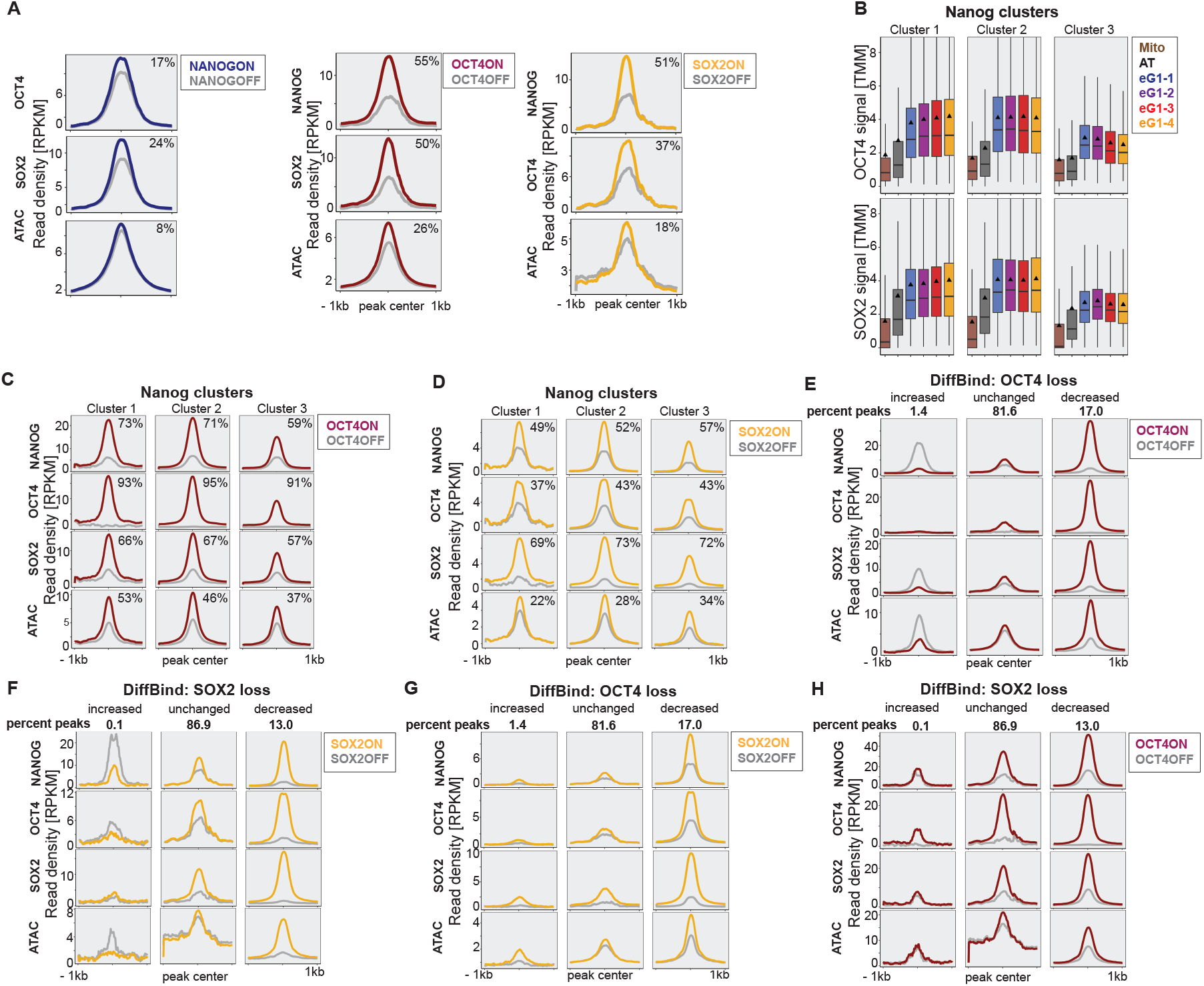
The interaction between OCT4 and SOX2 with NANOG is locus-dependent. (A) RPKM-normalized read density of OSN and chromatin accessibility in NANOG^cond^, OCT4^cond^, or SOX2^cond^ mESCs before and after 24 h doxycycline removal or treatment, or 26 h, respectively, centered around all OSN or merged peaks of the two TFs above (ATAC-seq). The percentage in the top right corner indicates the fold change between read enrichment in the presence or absence of the indicated TFs. (B) Boxplots of TMM-signal of OCT4 or SOX2 from mitosis to eG1-4 across all NANOG binding sites in Nanog clusters 1-3. Boxes: intervals between the 25th and 75th percentile. Horizontal lines: median; Triangles: mean; Error bars: 1.5-fold the interquartile range or the closest data point when the data point is outside this range. (C-D) RPKM-normalized read density of OSN and chromatin accessibility in OCT4^cond^ (C) or SOX2^cond^ mESCs (D) before and after doxycycline treatment, centered around all NANOG sites of clusters 1, 2, or 3. Percentage in the top right corner: fold change between read enrichment in the presence or absence of the indicated TFs. (E-H) RPKM-normalized read density of OSN and chromatin accessibility in OCT4^cond^ (E, H) or SOX2^cond^ mESCs (F, G) before and after doxycycline treatment, centered around all NANOG loci that increase, maintain, or decrease NANOG binding upon OCT4 (E, G) or SOX2 loss (F, H). Percentage of loci that increase, maintain, or decrease binding from the total number of sites is indicated on top.

**Figure S4 – related to Figure 5:**
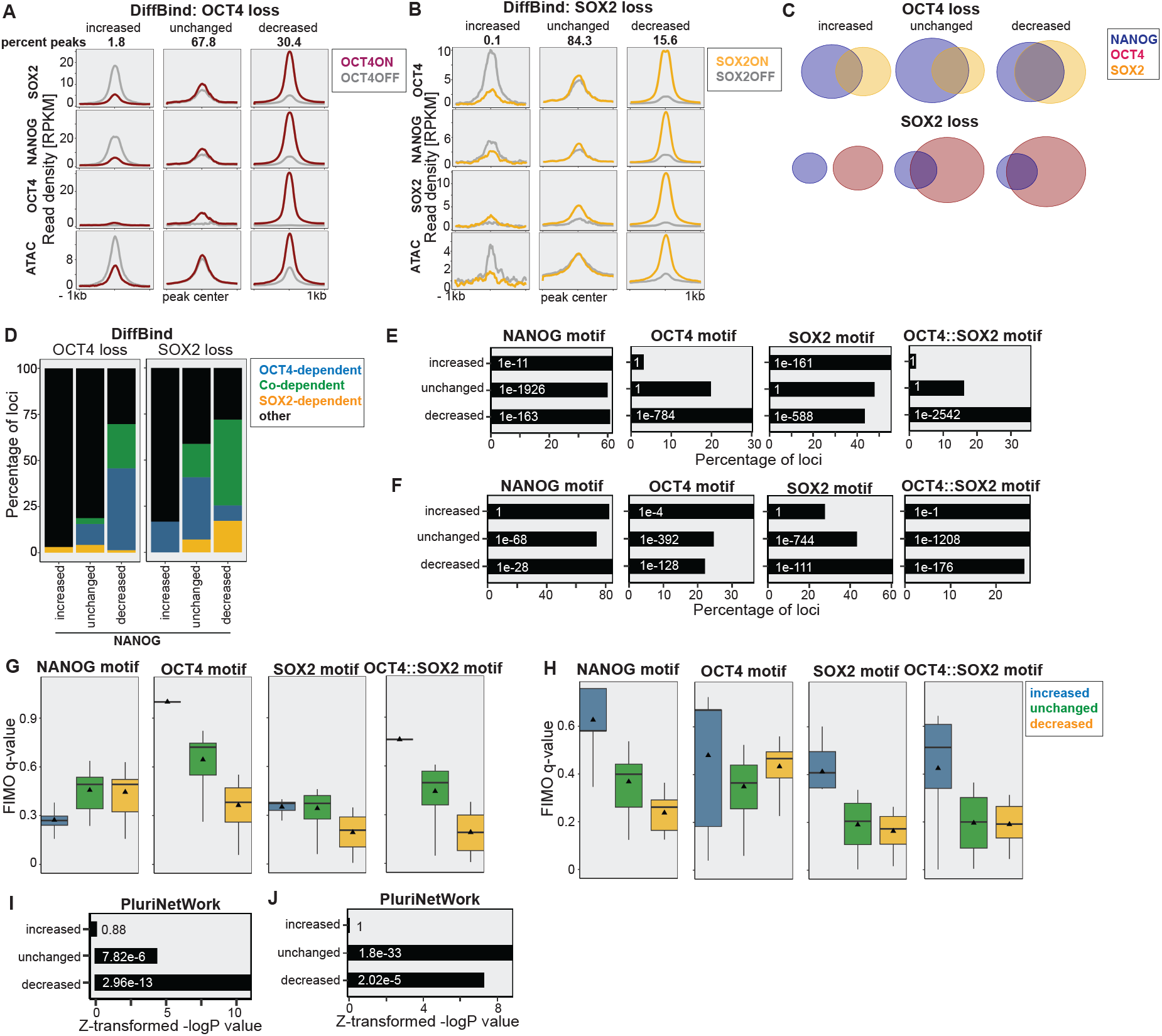
Additional characterization of significantly affected loci in the absence of OCT4 or SOX2. (A-B) RPKM-normalized read density of OSN and chromatin accessibility in OCT4^cond^ (A) or SOX2^cond^ mESCs (B) before and after 24 h doxycycline treatment, or 26 h, respectively, centered around all SOX2 or OCT4 loci that increase, maintain or decrease SOX2 or OCT4 binding. (C) Venn diagrams show the overlap between loci that significantly change NANOG or SOX2 binding upon OCT4 loss or NANOG and OCT4 binding upon SOX2 loss, respectively. (D) Overlap of the binding loci that significantly increase, maintain, or decrease NANOG in the absence of OCT4 or SOX2 with OCT4-, SOX2-, and Co-dependent loci. (E-F) NANOG, OCT4, SOX2, and OCT4::SOX2 motif enrichment and p-value (digits) at loci that significantly increase, maintain, or decrease NANOG in the absence of OCT4 (E) or SOX2 (F). (G-H) NANOG, OCT4, SOX2, and OCT4::SOX2 motif quality based on FIMO q-value at loci that significantly increase, maintain, or decrease NANOG binding in the absence of OCT4 (G) or SOX2 (H). Boxes: intervals between the 25th and 75th percentile. Horizontal lines: median; Triangles: mean; Error bars: 1.5-fold the interquartile range or the closest data point when the data point is outside this range. (I-J) Relative enrichment values plotted as Z-transformed -logP value and p-values (digits) of the GO terms PluriNetWork and Embryo development for loci that significantly increase, maintain, or decrease NANOG in the absence of OCT4 (I) or SOX2 (J).

### NANOG stabilizes OCT4 and SOX2 binding

While near-complete NANOG loss (Figure S5A) had only a mild effect on chromatin accessibility and on OCT4 and SOX2 binding (Figure 5A), we reasoned that NANOG may still strengthen OCT4 and SOX2 binding in naive PSCs. We found that 10.3 % and 12.9 % of loci only moderately decreased OCT4 or SOX2 binding in the absence of NANOG, respectively, and that a majority of these were common between for OCT4 and SOX2 (Figure 6A-B, Figure S5B, Table S3-4). Furthermore, the loci that increased OCT4 binding in the absence of NANOG decreased OCT4 binding in the absence of SOX2, whereas loci that increased SOX2 binding in the absence of NANOG remained unchanged in the absence of OCT4 (Figure 6C-D). Loci with increased OCT4 and SOX2 binding in the absence of NANOG also had a higher abundance of the SOX2 motif compared to those with decreased OCT4 and SOX2 binding (Figure S5C-D). This indicates that in the absence of NANOG, SOX2 redirects OCT4 to new loci and not the other way around. Loci that significantly decreased OCT4 or SOX2 binding upon NANOG loss were enriched for all four pluripotency motifs (Figure S5C-D), as well as for the GO terms pluripotency and embryo development (Figure S5E-F), consistent with the sustained self-renewal and increased susceptibility to differentiation resulting from NANOG depletion (Chambers et al., 2007).

**Figure 6.**
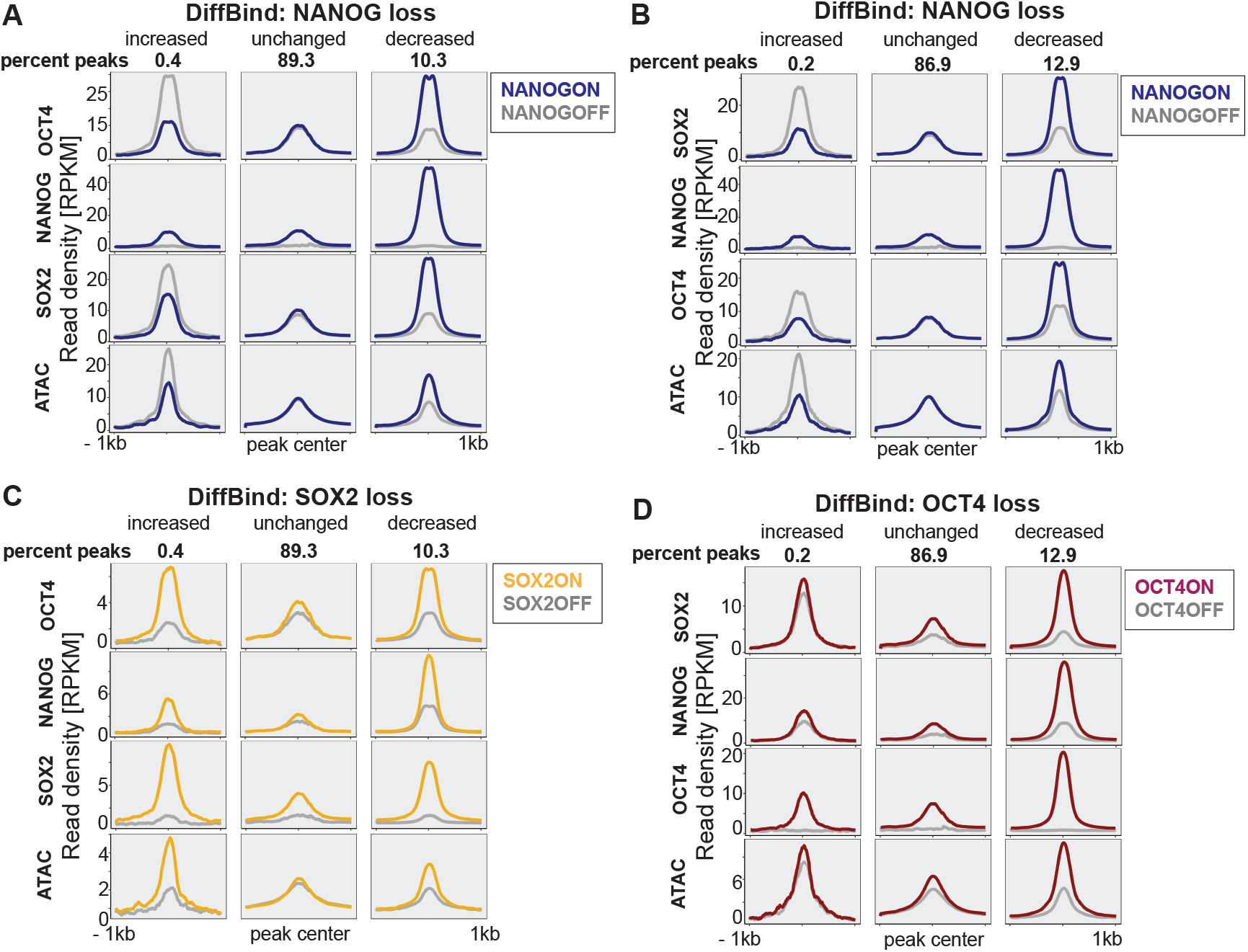
NANOG loss redistributes OCT4 and SOX2. (A-D) RPKM-normalized read density of OSN and chromatin accessibility in NANOG^cond^ (A, B), SOX2^cond^ (C), and OCT4^cond^ mESCs before and after 24 h doxycycline removal or treatment, or 26 h treatment, respectively, centered around all OCT4 (A, C) or SOX2 (B, D) loci that increase, maintain, or decrease NANOG binding.

**Figure S5 – related to Figure 6:**
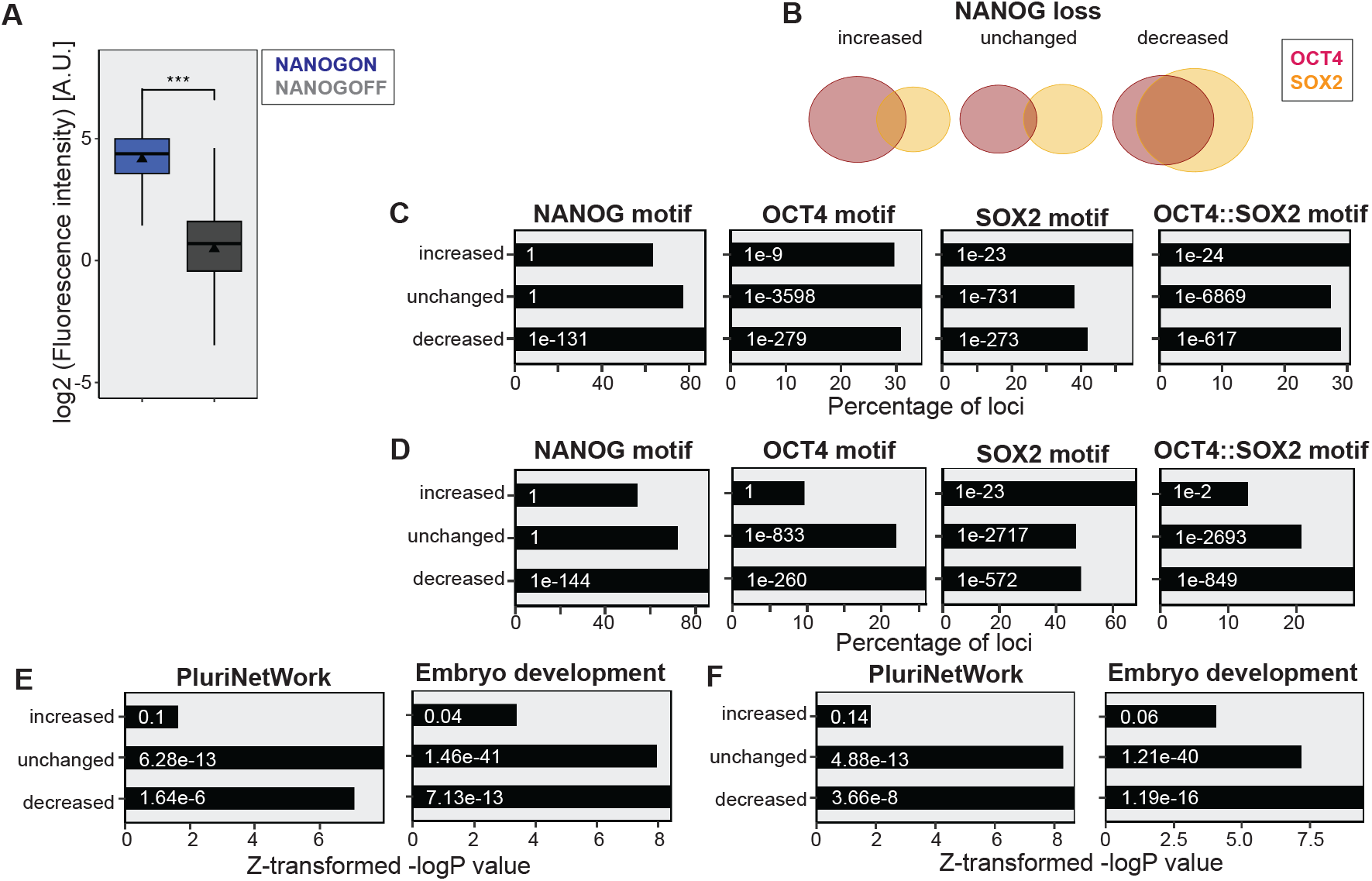
Additional characterization of significantly affected loci in the absence of NANOG. (A) Quantification of the immunofluorescence on NANOG before and after doxycycline removal for 24 h. Boxes: intervals between the 25th and 75th percentile. Horizontal lines: median; Triangles: mean; Error bars: 1.5-fold the interquartile range or the closest data point when the data point is outside this range. (B) Venn diagrams showing the overlap between loci that significantly change OCT4 or SOX2 binding upon NANOG loss. (C-D) NANOG, OCT4, SOX2, and OCT4::SOX2 motif enrichment and p-values (digits) at loci that significantly increase, maintain or decrease OCT4 (C) or SOX2 (D) binding in the absence of NANOG. (E-F) Relative enrichment values plotted as Z-transformed -logP value and p-values (digits) of GO terms PluriNetWork and Embryo development for loci that significantly increase, maintain, or decrease OCT4 (E) or SOX2 (F) in the absence of NANOG.

In summary, NANOG mainly stabilizes the binding of OCT4 and SOX2, and in its absence OCT4 is partially redirected by SOX2 to other loci.

### OCT4 becomes increasingly important to maintain NANOG and SOX2 at pluripotency loci during naïve pluripotency exit

Finally, we asked whether the capacity of SOX2 to redirect NANOG binding in the absence of OCT4 is strengthened during exit of naïve pluripotency, during which SOX2 and NANOG may start to display functions more related to differentiation. To do so, we took advantage of a published dataset in which OCT4 was depleted in PSCs cultured in serum + LIF (SL) (King and Klose, 2017). Upon loss of OCT4, NANOG significantly gained and lost binding at 25 % and 8 % of its binding sites (Figure 7A, Table S3), respectively (Figure 5E). Most loci were common between different medium conditions (Figure 7B, Table S4), and increased NANOG binding was accompanied by an increase in SOX2 binding and chromatin accessibility (Figure 7A). Many loci that increased NANOG binding also significantly increased SOX2 binding (Figure 7C, Table S4) and were highly enriched in SOX2 motifs (Figure 7D). During development of the early blastocyst (E3.5), NANOG and SOX2 co-occupy sites with GATA6 at pre-endoderm and epiblast cis-regulatory elements (Thompson et al., 2022). We asked whether this co-occupancy is fostered in the absence of OCT4. Loci that significantly increased binding upon OCT4 loss were enriched for the GATA6 motif (Figure 7D) and in embryo development GO terms, while they were depleted in pluripotency GO terms (Figure 7E). Furthermore, using published GATA6 ChIP-seq (Wamaitha et al., 2015) acquired 36 h after Gata6 expression in mESCs, we found that GATA6 was more enriched at loci that increased compared to loci that decreased NANOG binding in the absence of OCT4 (Figure 7F-G). Altogether, our results suggest that during progression from naive to primed pluripotency, OCT4 plays an increasingly important role in maintaining NANOG and SOX2 binding at pluripotency regulatory elements and preventing their binding to loci involved in fostering further developmental stages.

**Figure 7.**
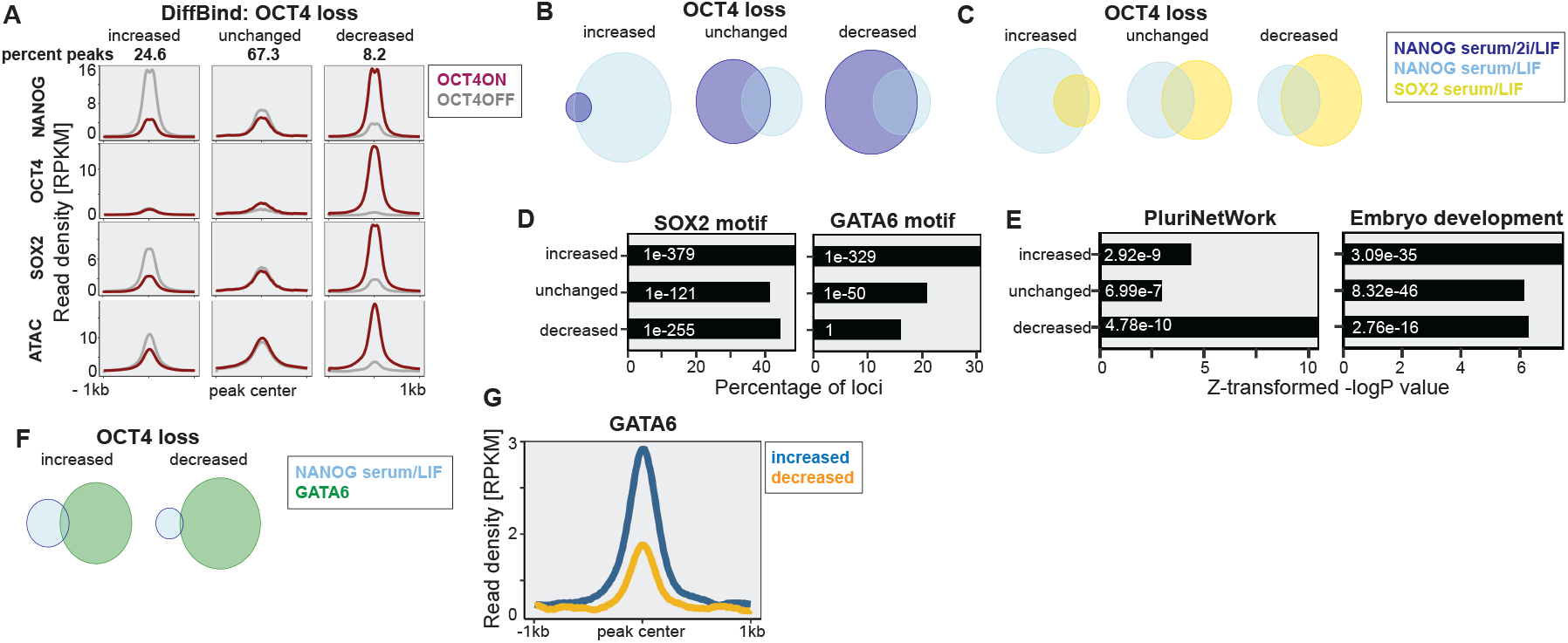
OCT4 is required to maintain NANOG at pluripotency regulatory elements in PSCs cultured in serum + LIF. (A) RPKM-normalized read density of OSN and chromatin accessibility in OCT4^cond^ mESCs grown in serum + LIF before and after 24 h doxycycline treatment centered around all NANOG loci that increase, maintain, or decrease NANOG binding. (B) Venn diagrams show the overlap between loci that significantly change NANOG binding upon OCT4 loss in serum/2i/LIF or serum/LIF. (C) Venn diagrams show the overlap between loci that significantly change NANOG or SOX2 binding upon OCT4 loss. (D) SOX2 and GATA6 motif enrichment and p-values (digits) at loci that significantly increase, maintain, or decrease NANOG in the absence of OCT4. (E) Relative enrichment values plotted as Z-transformed -logP value and p-values (digits) of GO terms PluriNetWork and Embryo development for loci that significantly increase, maintain, or decrease NANOG binding in the absence of OCT4. (F) Venn diagrams show the overlap between loci that significantly increase or decrease NANOG binding upon OCT4 loss with all GATA6 binding sites. (G) RPKM-normalized read density of the GATA6 signal centered around loci that increase or decrease NANOG binding in the absence of OCT4.

## Discussion

Here we describe a drug synchronization-free approach to dissect the orchestration of TF binding, chromatin accessibility and transcriptional activity during mitotic exit. We show that the pluripotency network is quickly consolidated during mitotic exit, while TF binding events and mRNAs involved in differentiation are rapidly turned off.

Several studies using nocodazole-release approaches have reported a major spike in transcriptional activity when cells exit mitosis (Chervova et al., 2023; C-S Hsiung et al., 2016; Palozola et al., 2017). Here in contrast, we report that transcription is progressively ramping up during eG1. While we cannot exclude that cell type-specific differences contribute to these discrepancies, this raises the possibility of unphysiological transcriptional reactivation upon nocodazole release.

The decrease in site-specific binding of most TFs during mitosis is well-documented, but it was so far unclear how they dynamically re-establish their occupancy landscape during mitotic exit. Using unsynchronized cells, we confirmed that OSN are mostly evicted from specific binding sites during mitosis, as reported before in nocodazole-treated cells (Deluz et al., 2016; Festuccia et al., 2019). The rapid re-initiation of their genome-wide site-specific binding at the anaphase-telophase transition is consistent with the re-establishment of promoter-enhancer loops in this time window (Pelham-Webb et al., 2021; Zhang et al., 2019). We also found that OSN exhibit markedly distinct temporal rebinding patterns to the accessible genome. The pioneer TF SOX2 first explores accessible regions before relocating to more closed sites. In contrast, OCT4 becomes progressively relocated to more accessible regions of the genome during eG1. Since OCT4 has a strong, genome-wide pioneering activity in PSCs (Friman et al., 2019; King and Klose, 2017) this suggests that OCT4 is the main driver in re-establishing full post-mitotic chromatin accessibility at its binding sites. Like OCT4, NANOG also first explores inaccessible sites before being relocated to more accessible regions. However, NANOG depletion had only a minor impact on chromatin accessibility, in line with its lack of pioneering activity in mouse PSCs.

We also demonstrate the distinct roles of OCT4 and SOX2 in guiding NANOG and each other’s binding. SOX2 can redirect NANOG or OCT4 to different genomic loci in the absence of OCT4 or NANOG, respectively. In contrast, OCT4 plays an invariable role in attracting SOX2 and NANOG towards pluripotency regulatory regions, independently of their respective expression. Our study thus reveals the diversity in which TFs adapt their binding landscape or recruit other TFs depending on the availability of their binding partners. Importantly, this also illuminates the temporal orchestration of TF binding during mitotic exit. In eG1-1, NANOG binding is biased towards differentiation-associated loci compared to later time points, with a higher dependence on the pioneering activities of SOX2 alone than OCT4 or both together. This suggests that SOX2 has a larger impact on NANOG binding than OCT4 early during mitotic exit, which could be partly mediated by its stronger physical interaction with NANOG (Gagliardi et al., 2013; Mistri et al., 2022). In contrast, OCT4 alone and in cooperation with SOX2 progressively shifts the NANOG binding landscape towards globally more accessible and pluripotency-related regions. Since the co-localization of OSN does not substantially vary during eG1, the temporal changes in NANOG binding related to OCT4 and SOX2 are likely to be mostly indirectly mediated by their pioneering activity. These findings shed light on a brief post-mitotic temporal window during which the pluripotent TF binding landscape is not fully re-established yet, which may be involved in the increased sensitivity to differentiation during the mitosis-G1 transition (Coronado et al., 2013; Halley-Stott et al., 2014; Pauklin and Vallier, 2013).

### Limitations of the study

Although the Cdt1-MD reporter system allowed us to efficiently distinguish between different temporal windows of eG1, the gate size had to be adjusted for each cell sorting experiment. Together with variability in ChIP-seq efficiency between each replicate this may contribute to the sawtooth aspect of the ChIP-seq signal in (Figure S2B). Thus, we decided to rely on TMM-normalized data for most of our quantitative analysis, focusing on relative, locus-specific changes in TF binding. However, although TMM allows us to overcome the technical limitation of sample preparation, we cannot rule out that normalization masks some true changes in global TF occupancy. Furthermore, our AT-enriched cell population was only enriched for 32% of anaphase-telophase cells, and thus the overall signal from this population is likely to not truly reflect the full extent of OSN occupancy.

## Acknowledgements

This study was funded by the Swiss National Science Foundation (grant# 310030_184782). We thank the whole Gene Expression Core Facility from EPFL, especially Bastien Mangeat, Elisa Cora, and Lionel Ponsonnet, for next-generation sequencing and library preparation of the RNA-seq samples. We thank the Flow Cytometry Core Facility at EPFL, especially Miguel Garcia, Valérie Glutz, Francesco Palumbo, and André Mozes (former member) for cell sorting and provision of flow cytometry equipment. We thank the Bioimaging and Optics Platform and Biomolecular Screening Facility at EPFL for assistance with microscopes, SCITAS for cluster computing, Pablo Navarro for sharing 44iN cells with us, Armelle Tollenaere for helping with the collection of mitotic cells and for critical reading of the manuscript, and Lucas Jamar for helping with the linear regression.

## Author contributions

Conceptualization, S.P: and D.S.; methodology, S.P. and D.S; investigation, S.P. and L.V.; formal analysis, S.P. and D.S.; resources, D.S; data curation, S.P.; writing – original draft, D.S and S.P.; writing – review & editing, S.P., L.V. and D.S; supervision, D.S.; funding acquisition, D.S.

**Video S1 – related to Figure 1: Video of mESCs stably expressing the Cdt1-MD reporter.**

Images were taken every 15 min for 76 timeframes at a magnification of 20x. mCherry: red; YPet: grey.

## STAR Methods

### Cell culture

The CGR8 (ECACC 07032901), ZHBTc4 (Niwa et al., 2000), 2TS22C (Masui et al., 2007), and 44iN mouse embryonic stem cells lines (Festuccia et al., 2012) were routinely cultured on 0.1% gelatin-coated (Sigma, G9391) 100mm Petri dishes at 37°C and 5% CO_2_ in GMEM (Sigma, G5154) supplemented with 10% inactivated ESC qualified fetal bovine serum (Gibco, 16141–079), 1% nonessential amino acids (Gibco,11140–050), 2 mM L-glutamine (Gibco, 25030–024), 2 mM sodium pyruvate (Sigma, S8636-100ML), 100µM 2-mercaptoethanol (Sigma, M3148), 1% penicillin and streptomycin (BioConcept, 4–01F00-H), leukemia inhibitory factor (made in-house), 3 µM CHIR99021 (Merck, 361559) and 0.8 µM PD184352 (Sigma, PZ0181). Cells were split every 2-3 days based on their confluency by trypsinization (Sigma, T4049). 44iN cells were kept in a medium containing 1 µg/ml doxycycline (Sigma, D3447). For ChIP-seq, ATAC-seq and RNA-seq experiments on the sorted cell populations of the Cdt1-MD cell line, cells were cultured for at least one week in a 1:1 mix of Neurobasal Medium (Gibco, 21103049) and DMEM/F-12 (Gibco, 21331020), supplemented with B-27 (Gibco, 17504044), N-2 (Gibco, 17502048), 2 mM L-glutamine (Gibco, 25030–024), 1% penicillin and streptomycin (BioConcept, 4–01F00-H), 100µM 2-mercaptoethanol (Sigma, 63689), 3 µM CHIR99021 (Merck, 361559), 1 µM PD184352 (Sigma, PZ0181), and leukemia inhibitory factor (in-house produced and validated for its capacity to maintain pluripotency). HEK 293T cells were cultured in DMEM with GlutaMAX (Gibco, 31966021) supplemented with 10% fetal bovine serum (Gibco, 10270106) and 1% penicillin and streptomycin (BioConcept, 4–01F00-H). To deplete OCT4 and SOX2 in the ZHBTc4 and 2TS22C cell lines, respectively, 10^6^ cells were seeded in a 10cm dish and the next day, ZHBTc4 and 2TS22C cells were treated with 1 µg/ml doxycycline for 24 h or 26 h, respectively, before fixation. To deplete NANOG from 44iN cells, 10^6^ cells were seeded in a 10cm dish in the presence of 1 µg/ml doxycycline, and the next day the medium was changed to leave cells in doxycycline-free medium for 24 h.

### Plasmid construction

The lentiviral vector pLV-PGK-YPet-MD was previously generated in the Suter laboratory (Friman et al., 2019). For the construction of pLV-EF1a-mCherry-hCdt1(1/100)(Cy-), we inserted hCdt1(1/100)(Cy-) (Invitrogen, GeneArt Gene Synthesis) in fusion to mCherry in a pLV-EF1a-mCherry/PGK-Hygromycin vector by in-fusion cloning (Takara Bio, 639648).

### Lentiviral vector production and stable cell line generation

HEK 293T cells were transfected with the psPAX2 (Addgene, 12260) envelope plasmid, the pMD2.G (Addgene, 12259) packaging plasmid (Dull et al., 1998), and pLV-PGK-YPet-MD or pLV-EF1a-mCherry-hCdt1(1/100)(Cy-) using calcium phosphate transfection (Suter et al., 2006). Two days after transfection, lentiviral vectors were concentrated by ultracentrifugation for 2 h at 20’000 g and 4°C. Subsequently, 50,000 mESCs plated in a 24-well plate in 1 ml of medium were transduced with 50 µl of concentrated pLV-EF1a-mCherry-hCdt1(1/100)(Cy-) lentiviral particles. mCherry-positive cells were selected with 200 µg/ml hygromycin B (Gibco, 10687010) for 10 days, and were subsequently transduced with pLV-PGK-YPet-MD viral particles. To generate a clonal cell line, single mCherry and YPet double-positive cells were sorted in individual wells of a 96-well plate and kept under selection with hygromycin. After clonal outgrowth, clones were further passages and one clone with high mCherry and YPet expression levels was selected as the final Cdt1-MD cell line.

### Sorting of different cell cycle phases

Cells were stained in suspension with 40.5 µM Hoechst 33342 (Thermo Fisher Scientific, H3570), and incubated for 20 min on a rotating platform at room temperature. The tube was filled with PBS + 1% FBS and spun down at 600 g for 5 min. For the sorting of different temporal bins in eG1, cells were fixed as described in the section “ChIP-seq”, resuspended in cold PBS + 1% FBS and sorted at 4°C based on their mCherry and YPet intensity in G1. Flow cytometry and FACS were performed on a LSR Fortessa (BD Biosciences) and on a FASCAria Fusion (BD Biosciences), respectively. The gates were adjusted to obtain the same number of cells for each temporal bin. To obtain mitotic or telophase-anaphase enriched cells, cells were spun down at 450 g for 5 min at 4°C following fixation and Hoechst staining. Subsequently, cells were resuspended in 225 µl PBS and permeabilized by adding 4.5 ml cold 70% EtOH (Fisher Scientific, 10342652) in PBS under vortex agitation and incubated for 15 min on ice with frequent inversion. Cells were then washed twice by resuspending them in PBS + 1% FBS, followed by centrifugation at 450 g for 5 min at 4°C. Following the second washing step, cells were resuspended in PBS + 10% FBS ES and the primary anti-H3S10p antibody (Millipore, 05-1336) was added at a dilution of 1:1000 and incubated for 90 min at 4°C in the dark. Cells were then washed twice with PBS + 1% FBS and centrifuged at 450 g for 5 min at 4°C. Subsequently, cells were resuspended in PBS + 10% FBS, the secondary antibody donkey anti-mouse IgG-AF647 (Invitrogen, A31571) was added at a dilution of 1:80, and cells were incubated for 45 min at 4°C in the dark. Finally, cells were washed twice with PBS + 1% FBS, centrifuged at 450 g for 5 min at 4°C, and resuspended in PBS + 1% FBS for FACS. The anaphase-telophase-enriched population was selected by gating for H3S10p+, mCherry-high, YPet-intermediate, and Hoechst-high cells. Mitotic cells were selected by gating for Hoechst-high and H3S10p+ cells. The data was analyzed using the FlowJo software (version 10.8.1). To obtain 1 million mitotic cells for OCT4 ChIP-seq from unsynchronized populations (Figure S2A), 580 Mio of CGR8 cells were plated into 145 T75 flasks. The next day, we used mitotic shake-off followed by sorting of H3S10p+ cells using FACS to purify mitotic cells.

### ATAC-seq

ATAC-seq experiments were performed in biological duplicates for all sorted and unsorted cell populations. The experimental procedures were followed as previously described (Buenrostro et al., 2013). Briefly, 50,000 cells were sorted or collected after trypsinization. Cells were then pelleted at 800g for 5min at 4°C, washed once with cold PBS, and kept on ice. Subsequently, cells were resuspended in 50 µl cold lysis buffer (10 mM Tris-HCl pH 7.4, 10 mM NaCl, 3 mM MgCl2, 0.1% NP-40), and centrifuged at 800g for 10min at 4°C. For the transposition reaction, pelleted nuclei were resuspended in 50 µl of 0.5 µM Tn5 (in-house produced according to (Chen et al., 2017)) in TAPS-DMF buffer (10 mM TAPS-NaOH, 5 mM Mgcl2, 10% DMF) and incubated at 37°C for 30 min. During the reaction, the tubes were flicked gently several times, and DNA was immediately purified using the DNA Clean & Concentrator kit (Zymo Research, D4003) with a binding buffer to sample ratio of 5:1. Transposed DNA was diluted in 10 µl DNA elution buffer, and amplified in a solution containing 0.5 mM Ad1.1 universal primer, 0.5 mM of Ad2.X indexing primer, 0.6X SYBR Green I (Lonza, 59513) and 1X NEBNext High-Fidelity 2X PCR Master Mix (NEB, M0541L) at 72°C for 5 min, 98°C for 30 s, and 5 cycles of 98°C for 10 s, 63°C for 30 s, and 72°C for 1 min. The PCR reaction was monitored to stop the amplification before saturation by sampling 10 µl of amplified DNA and analyzing it by qPCR (Applied Biosystems, 7900HT). The number of cycles to add was determined by the time at which the fluorescence was still in the rising phase. The remaining DNA was further amplified at 98°C for 30 s, and 4-7 cycles of 98°C for 10 s, 63°C for 30 s, and 72°C for 1 min. Next, the library was purified using the DNA Clean & Concentrator kit (Zymo Research, D4003) with a binding buffer-to-sample ratio of 5:1. The purified library was further size-selected using a 0.55X and 1.2X ratio of AMPure XP beads (Beckman Coulter, A63880). Library concentrations were determined by Qubit, analyzed on a fragment analyzer, and sequenced on an Illumina NextSeq 500 using 75 nucleotide read-length paired-end sequencing.

### ChIP-seq

ChIP-seq experiments for Cdt1-MD cells in eG1-1 to eG1-4 on OCT4, and eG1-1 and eG1-3 for SOX2 were acquired in biological triplicates, the experiments on 50,000 cells in one biological replicate, and for the remaining samples biological duplicates were acquired. Briefly, cells were collected after trypsinization and fixed with 2 mM DSG (Thermo Scientific, 20593) in PBS for 50 min at room temperature on a rotating platform. Subsequently, cells were pelleted for 5 min at 600 g, and fixed for 10 min with 1 % formaldehyde (Thermo Scientific, PIER28906) on a rotating platform. The reaction was stopped using 200 mM Tris-HCl pH 8.0 (PanReac AppliChem, A4577) for 10 min on a rotating platform. Tubes were filled with 1 % FBS in PBS and cells were spun for 5 min at 600 g and 4°C. Fixed cells were either directly further processed or stained for cell sorting, as described in the “Sorting of different cell cycle phases” section. Nuclei were extracted by resuspending the cell pellet twice in LB1 (50 mM HEPES-KOH pH 7.4, 140 mM NaCl, 1 mM EDTA, 0.5 mM EGTA, 10% Glycerol, 0.5% NP-40 (Thermo Scientific, 85124), 0.25% Triton X-100), incubated for 10 min at 4°C and with gentle shaking at 100 rpm, and pelleted for 5 min at 4°C and 1700 g. The nuclei were then resuspended in LB2 (10 mM Tris-HCl pH 8.0, 200 mM NaCl, 1 mM EDTA, 0.5 mM EGTA), incubated for 10 min at 4°C and with gentle shaking at 100 rpm, and spun down for 5 min at 4°C and 1700 g. To wash nuclei, SDS shearing buffer (10 mM Tris-HCl pH 8.0, 1 mM EDTA, 0.15% SDS) was added twice without disturbing the pellet, the tube was rolled to rinse the walls and spun for 5 min at 4°C and 1700 g. All buffers were supplemented with a 1:100 dilution of protease inhibitor cocktail (Sigma, P8340). Chromatin was sheared for 20 min at 5% duty, 140 W, 200 cycles, or for samples with 50,000 cells for 10 min, 10% duty, 75 W, 200 cycles in a Covaris E220 focused ultrasonicator. To remove the insoluble material, chromatin was spun for 10,000 g for 5 min at 4°C and the supernatant was transferred to a new tube. As total input control and to evaluate the sonication efficiency, a few microliters of chromatin were incubated with 1X TE buffer pH 8.0 (PanReac AppliChem, A0386) and 10 ng/µl RNAse A (Qiagen, 19101) at 37°C for 30 min. Proteinase K (Qiagen, 19131) was added at 400 ng/µl and the reaction was incubated for another 30 min at 55°C and shaking at 1100 rpm. Finally, 200 mM NaCl (Sigma, 59222C) was added to reverse the crosslinks at 65°C for 16 h and shaking at 1100 rpm. The chromatin was purified using a MinElute PCR purification kit (Qiagen, 28004) and quality-checked on a 1 % agarose gel (Invitrogen, 16500500). Next steps were performed using the ChIP-IT High Sensitivity kit (Active motif, 53040) with a few modifications highlighted below. For immunoprecipitation of samples made from at least 1 million cells, 3.5 µg chromatin were used unless stated otherwise, and for samples made from 50,000 cells, all extracted chromatin was used. The following antibodies were used: anti-OCT4A (Cell Signaling Technology, 5677S) at 3.1 µl per 3.5 µg chromatin, anti-SOX2 (Cell Signaling Technology, 23064) at 3.1 µl per 3.5 µg chromatin, and anti-NANOG (Cell Signaling Technology, 8822) at 1.55 µl per 3.5 µg chromatin. To obtain quantitative ChIP-seq results, drosophila spike-in chromatin (Active motif, 53083) and spike-in antibody (Active motif, 61686) were used according to the manufacturer’s recommendations. Before binding the immunoprecipitated chromatin to the beads, the beads were washed once with ChIP buffer. For cross-link reversal, 2 µl proteinase K was added, the reaction was incubated at 55°C for 30 min and the temperature increased to 65°C and shaking at 1100 rpm for 16 h. Sequencing libraries were prepared using the NEBNext Ultra II DNA Library Prep Kit (NEB, E7645L), and sequenced on a NextSeq 500 using 75-nucleotide read length paired-end sequencing.

### Nuclear RNA-seq

Nuclear RNA-seq was done in biological duplicates on the four sorted temporal bins of eG1. Upon sorting, cells were spun down at 600 g for 5 min at 4°C, and nuclei were isolated following the low cell input nuclei isolation protocol of the 10x protocol for cell lines and PBMCs (https://assets.ctfassets.net/an68im79xiti/6t5iwATCRaHB4VWOJm2Vgc/bdfd23cdc1d0a321487c8b231a448103/CG000365_DemonstratedProtocol_NucleiIsolation_ATAC_GEX_Sequencing_RevB.pdf). Briefly, cells were resuspended in 500 µl lysis buffer composed of 10 mM Tris pH 7.4, 10 mM NaCl, 3 mM MgCl2, 1% BSA (Merck, B9000S), 0.1% Tween (Fisher Scientific, 10113103), 1 mM DTT (Merck, 646563), 0.5U/µl Rnase inhibitor (Merck, 3335402001), 0.1% NP-40, and 0.01% Digitonin (Invitrogen, BN2006). After incubation for 5 min on ice, 500 µl wash buffer was added (same as lysis buffer but without NP-40 and digitonin) and nuclei were spun down at 450 g for 5 min at 4°C. The washing step was repeated for a total of three times. RNA extraction was done using the Rneasy Plus Micro Kit (Qiagen, 74034) with the following adjustments. Nuclei were resuspended in 350 µl lysis buffer comprising 40 mM DTT. Upon gDNA elimination, the samples were washed with ethanol and transferred to the RNeasy MinElute column. Next, 350 µl of RW1 buffer was added, and samples were spun at 8000 g for 30 s. The flow-through was discarded and a mix of 70 µl buffer RDD with 10 µl DNAse (Qiagen, 79254) was added to the columns. 15 minutes later, 350 µl buffer RW1 was added to the columns and the manufacturer’s protocol was followed. Sequencing libraries were prepared using the NEBNext Ultra II Directional RNA Library Prep Kit for Illumina (NEB, E7760S), and sequenced on a NextSeq 500 using 75-nucleotide read length paired-end sequencing.

### Immunofluorescence microscopy

Cells were plated in a 96-well plate pre-coated for 2 h at 37°C with 1:10 diluted Biolaminin (BioLamina, LN511-0202) in DPBS (Gibco, 14040091). The next day, 1 µg/ml doxycycline was added to ZHBTC4 or 2TS22C cells or removed from 44iN cells to deplete the corresponding TF for 24 h (ZHBTC4 and 44iN cells) or 26 h (2TS22C cells). Cells were then fixed with 2% formaldehyde in DPBS for 30 min at room temperature, washed with DPBS, and permeabilized with 0.5 % Triton X-100 in DPBS for 30 min at room temperature. After two washes with DPBS, cells were blocked for 30 min at room temperature with 1 % BSA in DPBS. The anti-Nanog antibody (CST, 8822) was added at 1:1000 dilution in a blocking solution, and the plate was incubated overnight at 4°C. After two washes with DPBS, the secondary anti-rabbit IgG AF647 antibody (Invitrogen, A-21443) was added at a dilution of 1:1000 in blocking solution for 1 h at room temperature in the dark. Cells were washed twice with 0.1 % Tween 20 in DPBS, then twice with DPBS, and Fluoromount G with DAPI (SouthernBiotech, 0100-20) was added and incubated for 10 min before imaging. The plate was imaged on an IN Cell Analyzer 2200 (GE Healthcare). Images were analyzed using CellProfiler (Carpenter et al., 2006). Nuclei were identified by area shape and size on the DAPI channel, and the integrated fluorescence intensity in the NANOG channel was measured in cell nuclei.

### Time-lapse imaging

Cdt1-MD cells were plated on a 96-well plate coated for 1h at 37°C with StemAdhere (Primorigen Biosciences, S2071) diluted 1:25 in DPBS. The medium was then exchanged for phenol red-free FluoroBrite DMEM (Gibco, A1896701) supplemented with 10% inactivated ESC qualified fetal bovine serum (Gibco, 16141–079), 1% nonessential amino acids (Gibco,11140–050), 2 mM L-glutamine (Gibco, 25030–024), 2 mM sodium pyruvate (Sigma, S8636-100ML), 100µM 2-mercaptoethanol (Sigma, M3148), 1% penicillin and streptomycin (BioConcept, 4–01F00-H), leukemia inhibitory factor (made in-house), 3 µM CHIR99021 (Merck, 361559) and 0.8 µM PD184352 (Sigma, PZ0181), and cells were imaged on an Operetta CLS (PerkinElmer) for 16 h every 15 min. To quantify YPet and mCherry signals, images were background-subtracted in Fiji (Schindelin et al., 2012) using a rolling ball radius of 50 pixels. The mean cell and background signal were tracked manually by maintaining the size of the region of interest over time. Signal measurements were started from the last timepoint before cell division, and one sister cell was followed for 5 h (21 frames). This captures the whole G1 phase in our Cdt1-MD cell line, as we determined by the average cell cycle length and the percentage of G1 cells. To obtain the integrated fluorescence intensity of individual cells, the background was subtracted from the mean fluorescence intensity multiplied by the area of the cell.

### Calculation of the post-cytokinesis time in the different temporal bins of eG1

To obtain the post-cytokinesis time of the eG1 bins, we extracted all measurements of the time-lapse imaging that were measured between timepoint 0 and 2 h since these correspond to the early timepoints of G1 based on the average plots of mCherry and YPet intensity and flow cytometry. Subsequently, we used scikit-learn (Buitinck et al., 2013) to model a linear regression line to cluster the data points into four equal-sized clusters with the most equal distribution in 2D. We extracted the data points and plotted them using ggplot (Wickham, 2009).

### Sequencing data analysis

Sequencing libraries were aligned to the *Mus musculus* mm10 genome (GRCm38 release M25) and to the *Drosophila Melanogaster* genome (BDGP6 release 28) using STAR 2.7.0e (Dobin et al., 2013). Duplicated reads were removed with Picard (Broad Institute), and indexed using SAMTools 1.10 (Li et al., 2009). ChIP-seq samples were spike-in-normalized based on the sample with the lowest abundance of *Drosophila* reads (Bonhoure et al., 2014; Orlando et al., 2014) using SAMTools. For replicates, bam files were merged using SAMTools, and BigWig files were computed using the deepTools 3.5.1 (Ramírez et al., 2016) function bamCoverage with the setting “--normalizeUsing RPKM”. Peaks were called using MACS 2.2.4 (Zhang et al., 2008) with settings “-f BAMPE -g mm”, and merged using awk. Blacklisted peaks (Dunham et al., 2012) were removed with BEDTools (Quinlan and Hall, 2010) using the setting “-v”.

The deepTools function computeMatrix with the setting “reference-point” was used to compute a matrix containing a score per genomic region. This matrix file was subsequently used together with the deepTools function plotHeatmap, and for average line plots with custom R code and the package ggplot2 (Wickham, 2009). The deepTools function multiBigWigSummary with the setting “BED-file” was used to calculate read counts per peak region and used together with ggplot2 to generate violin or box plots. BEDOPS (Neph et al., 2012) and BEDTools were used to merge and intersect different peak files, respectively. The deepTools function bigwigCompare with the setting “--operation subtract” was used to subtract BigWig files, or to obtain the quotient between two BigWig files using the setting “--operation ratio”. To upscale the mitotic and anaphase-telophase enriched samples, BigWig files were multiplied by the ratio between the 1 Mio and 50,000 cell sample of the corresponding eG1-1 files using IGB (Freese et al., 2016) and the resulting bedgraph files merged using the Unix command “cat”. The bedgraph files were converted to BigWig files using bedGraphToBigWig (Kent et al., 2010). To plot the subtracted tracks and multiplied BigWig files against the peaks, peaks were trimmed to 200 bp around the peak center using the HOMER2 function annotatePeaks.pl with the setting “-size 200” and the commands awk and tail. This trimming was performed to eliminate variability in peak signals emerging from differences in fragment sizes between samples. To identify the features of the OSN binding sites, target peaks were overlapped with ESC_J1 enhancer peaks from EnhancerAtlas (Gao and Qian, 2020) using BEDTools. For promoters, all TSS from the mm10 genome were extracted and enlarged by 1 kb upstream and 500 bp downstream, and also crossed with the target peaks using BEDTools. The remaining peaks were classified as other. TF clusters were obtained by calculating the log2fold change between different temporal bins of eG1 and eG1-1, and using the R function pheatmap with the settings “clustering_distance_rows = “euclidean”, kmeans_k = 3”. For the comparative analysis of TF binding in the presence or absence of another TF, DiffBind (Stark and Brown, 2011) was used with settings for dba.count “minOverlap = 2”, and only peaks with an FDR < 0.05 were kept. To obtain intronic or exonic reads from the total RNA-seq reads, we used SAMTools view. In the clusters of the RNA-seq data (Figure 2C), we only considered genes with at least one value over 0 for the intronic or exonic reads. However, for intron and exon plots (Figure 2B), all exons or all introns, respectively, were used.

### Motif and gene ontology analysis

For motif search, peak files were subjected to the HOMER2 (Heinz et al., 2010) function findMotifsGenome.pl with the setting “-size given”. We plotted the known motif and p-value of the target regions, and the background which is the mean of the estimated background frequency in all groups. Motif strength was calculated using FIMO (Grant et al., 2011) with the setting “--thresh 0.001” and motifs UN0383.1, MA1115.1, MA0142.1, and MA0143.1 were downloaded from JASPAR (Sandelin et al., 2004). For GO enrichment, the closest TSS to each peak was used for gene annotation either by using the R package ClusterProfiler (Wu et al., 2021; Yu et al., 2012) or for a functional analysis of enhancer-gene interactions from EnhancerAtlas by overlapping ESC_J1 peaks with the target peaks. The enrichment was calculated with findGO.pl from HOMER2 with the setting “mouse”, and plotted together with the Z-transformed -logP values.

### TMM normalization

For TMM normalization (Robinson and Oshlack, 2010), a peak file containing a list of all consensus peaks of OCT4, SOX2, and NANOG was generated using BEDOPS with the setting “-m”. The BED file was subsequently converted into the SAF format using awk, and a count matrix containing the information of all BAM files of interest was generated using featureCounts (Liao et al., 2014). Normalization factors were calculated using edgeR (Robinson et al., 2009), multiplied by the total library size, and divided by 10^6^ to scale to reads per million. Normalized BigWig files were computed using the tool bamCoverage of deepTools 3.5.1 with the settings “--scaleFactor” and “-bs 1”.

### Published datasets

Published data were downloaded from GEO (Edgar et al., 2002) and processed as described in the section “Data analysis for sequencing data”. ATAC-seq and ChIP-seq data from ZHBTc4 cells grown in serum/LIF conditions (GSE87822) (King and Klose, 2017), from 2TS22C cells (GSE134680) (Friman et al., 2019), and GATA6 ChIP-seq (GSE69323) (Wamaitha et al., 2015) were taken from previous studies. OCT4-, SOX2- and Co-dependent regions are coming from a previous study (Friman et al., 2019).

